# Decoding imagined speech reveals speech planning and production mechanisms

**DOI:** 10.1101/2022.05.30.494046

**Authors:** Joan Orpella, Francesco Mantegna, M. Florencia Assaneo, David Poeppel

## Abstract

Speech imagery (the ability to generate internally quasi-perceptual experiences of speech) is a fundamental ability linked to cognitive functions such as inner speech, phonological working memory, and predictive processing. Speech imagery is also considered an ideal tool to test theories of overt speech. The study of speech imagery is challenging, primarily because of the absence of overt behavioral output as well as the difficulty in temporally aligning imagery events across trials and individuals. We used magnetoencephalography (MEG) paired with temporal-generalization-based neural decoding and a simple behavioral protocol to determine the processing stages underlying speech imagery. We monitored participants’ lip and jaw micromovements during mental imagery of syllable production using electromyography. Decoding participants’ imagined syllables revealed a sequence of task-elicited representations. Importantly, participants’ micromovements did not discriminate between syllables. The decoded sequence of neuronal patterns maps well onto the predictions of current computational models of overt speech motor control and provides evidence for hypothesized internal and external feedback loops for speech planning and production, respectively. Additionally, the results expose the compressed nature of representations during planning which contrasts with the natural rate at which internal productions unfold. We conjecture that the same sequence underlies the motor-based generation of sensory predictions that modulate speech perception as well as the hypothesized articulatory loop of phonological working memory. The results underscore the potential of speech imagery, based on new experimental approaches and analytical methods, and further pave the way for successful non-invasive brain-computer interfaces.

## Introduction

Mental imagery of speech is the internally generated, quasi-perceptual experience of our own or others’ speech. Research on speech imagery has a long history in the sciences and philosophy^1–5^, for a variety of reasons. For one, speech imagery is integral to different cognitive functions, such as inner speech and phonological working memory, that play an important role in learning, problem-solving, and, more generally, development^6,7^. Speech imagery is also considered an informative model for overt speech^7–10^. As such, it has been employed to test aspects of speech planning, production, and motor control, which can be difficult to investigate with overt speech^8,9,11–15^. Mental imagery (of speech, or otherwise) is moreover a paradigmatic example of the generation of sensory predictions^8,16,17^. In speech, the hypothesis is that ‘the little voice in our head’ results from an internal prediction of the sensory consequences of planned motor commands^18^, that is, from a form of internal emulation^19^. These predictions can be used to anticipate sensory inputs, such as other’s speech, which facilitates comprehension^8^. This makes speech imagery an innovative tool to test predictive processing theories^20^, such as predictive coding^21,22^, Bayesian inference^23^, and associative learning^24–28^. Speech imagery is also clinically relevant. Imbalances between sensory predictions and feedback are thought to underlie disorders such as schizophrenia, autism, and stuttering^6,7,29^. Moreover, advances in the decoding of speech imagery are potentially life-changing for individuals that have lost the ability to speak due to stroke or illness^30–34^.

Despite its potential as a research and clinical tool, and despite the success of mental imagery research in other domains (e.g., limb motor control^35^, vision^36,37^), speech imagery remains poorly characterized. This is due to methodological challenges^6,38^ (e.g., the lack of comparative research, the lack of behavioral output, the potential timing misalignment across experimental trials and participants) and the absence of suitable paradigms and analytical approaches. Nevertheless, researchers have used speech imagery in creative ways, for example to quantify specific aspects of speech motor control (e.g., feedback prediction errors^13^), but the evidence remains indirect and incomplete. For instance, the experimental manipulation yielding speaking-induced suppression and its effects on perception can only *imply* the existence of so-called forward models and the precise sensory predictions emanating from planned speech^13,39,40^. The same methodological challenges permeate the attempts to decode speech imagery with time-resolved methods^38^, which have only recently begun to produce promising results, albeit restricted to state-of-the-art invasive (intracranial) recordings and complex analysis pipelines (e.g., ^41^).

We capitalized on the balance between temporal and spatial resolution afforded by the non-invasive method magnetoencephalography (MEG), paired with a deceptively simple speech imagery task (**Fig 1**) and a powerful decoding approach^42,43^ to determine the sequence of neural processes underlying speech imagery. In short, we decoded participants’ imagined speech as it unfolds.

**Fig 1.**
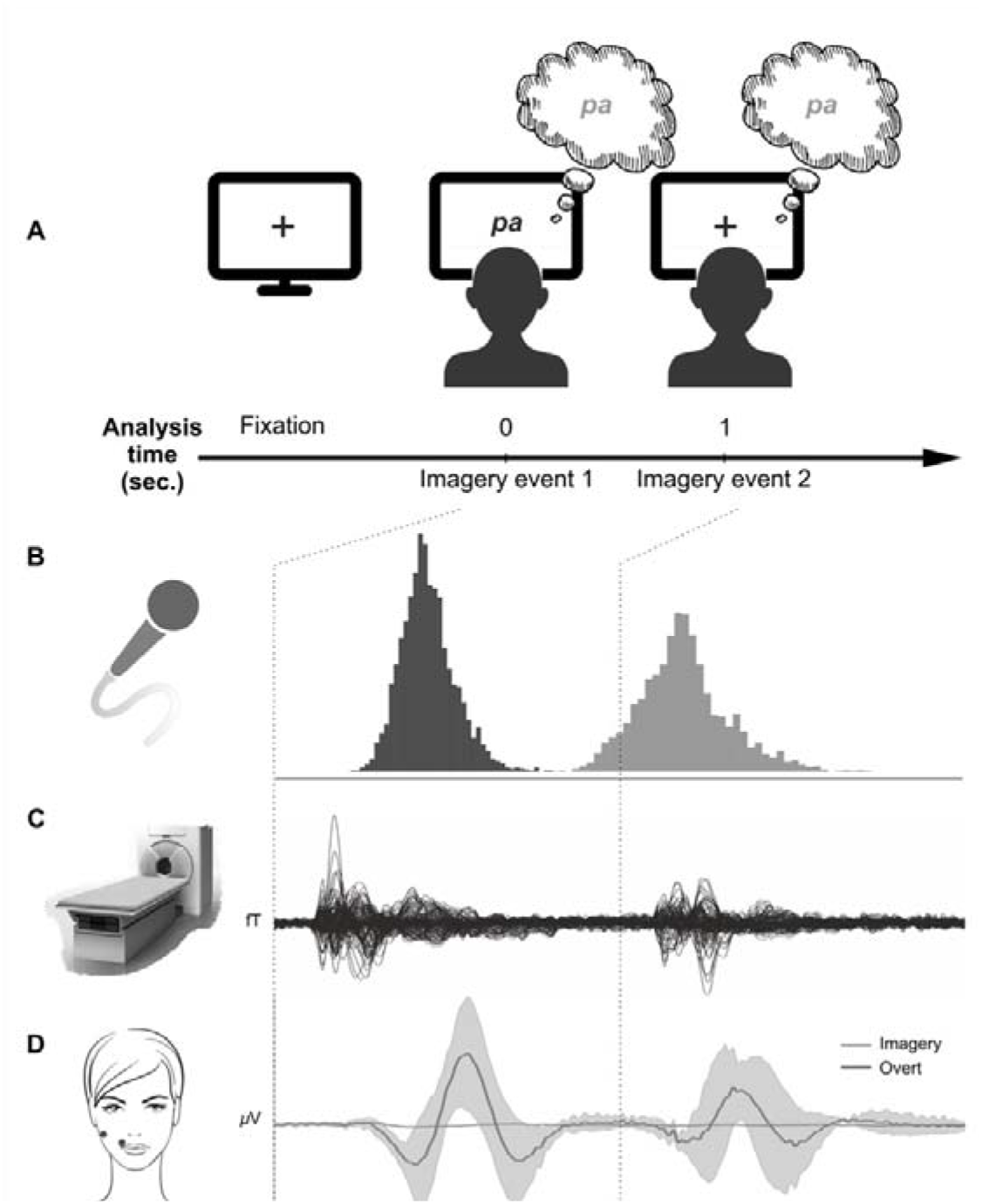
Experimental protocol. **A. Task.** Each trial began with a fixation cross of variable duration (1-1.5 sec) in the center of the screen. One of three syllables (e.g., /pa/, /ta/, /ka/) was then presented and remained on the screen for 1 second. Syllable presentation was followed by another fixation cross, lasting 2.5 seconds, after which the next trial began. The experiment comprised two conditions, *Imagery* and *Reading*, each with 4 blocks of 120 trials (40 presentations of each syllable per block), counterbalanced between participants. The total number of trials was 960 (320 per syllable). Syllable presentation was fully randomized within each block. In the *Imagery* blocks, participants were instructed to imagine producing the visually presented syllable as soon as possible after presentation (*event 1*) and a second time after second fixation cross presentation (*event 2*). In the *Reading* blocks, participants were instructed to fixate the center of the screen. Prior to the MEG session, participants received a training session: the experimenter explained the task and the desired type of imagery (i.e., imagining producing vs. hearing). Participants were also asked to complete a full block using *overt* productions of the syllables. At this time only, participants were given feedback regarding the timings of each production. This was critical to ensuring some minimal degree of temporal alignment within participant, as well as consensus across the cohort. Additionally, participants were given a link to an online version of the task (available here) to practice in their own time. **B. Estimating expected time of imagery**. On the day of MEG acquisition, each participant completed a minimum of 1 practice block (overt productions), which were recorded for subsequent analysis. Here all participants’ syllable onset distributions for events 1 and 2 are shown. The medians of these distributions were important for reference, to provide temporal boundaries for the times when imagery was to be expected. (See Fig S1 for additional data.) **C. Grand average MEG data for a participant’s *Imagery* trials**. Participants’ neural activity was recorded during both *Imagery* and *Reading* trials. These data (157 channels) were decoded for each participant. **D. Average electromyographic (EMG) data for one participant’s *Imagery* trials and for the same syllables spoken aloud**. To monitor participants’ potential micromovements during *Imagery*, we recorded muscle activity from the upper lip (lower dot) and jaw (upper dot) using a MEG-compatible EMG system. The figure exemplifies the magnitude of expected micromovements during imagery which, critically, did not differ between the imagined syllables (see Fig S2 – S4 for full analysis).

We recorded MEG signals from participants while they imagined internally producing isolated syllables (/pa/, /ta/, and /ka/) prompted on the screen on each trial. We used syllables as the targets of speech imagery given recent evidence for syllable-size ‘chunks’ as fundamental units for speech perception and production (^44^ for a review). First, we evaluated the extent to which these signals contained information over and above a *Reading* condition involving the passive viewing of the same syllables, and thus identical to *Imagery* except for the instruction to internally produce the prompted syllable. Having established robust differences between the two conditions, we next asked whether *Imagery* trials contained decodable content regarding the specific syllables imagined, that is, whether every syllable that participants imagine can be decoded from their neural data. Based on the decoding pattern, we then sought to characterize the entire genesis and development of the imagined speech events, a sequence which has so far remained elusive. We examined the dynamics of this sequence in order to deepen our understanding of how inner speech and sensory predictions are generated and ideally to adjudicate between current prevailing models of speech production, notably state feedback control (SFC)^8,9,23,45,46^ and DIVA^47^. Next, we examined the question of internal and external feedback loops for internal speech planning and production by assessing the time courses of auditory and motor areas during imagery. Lastly, we validated and extended the decoding results with data from a new cohort imagining a different set of syllables (/ta/, /tu/, and /ti/).

## Results

We recorded MEG signals from 21 participants (15 women; mean age = 28.19; *std* = 6.57) while they imagined producing one of three syllables (/pa/, /ta/, and /ka/). On every trial, participants were required to internally produce a given syllable as soon as it appeared on the screen (event 1) and a second time upon appearance of a fixation cross 1 second later (event 2) (**Fig 1A**). Electromyographic (EMG) data from the upper lip and jaw (**Fig 1D**) was acquired to measure any micromovements participants make during imagery. Although we expected micromovements during *Imagery*^48^, in line with previous research^7^, we did not expect these data to discriminate between different syllables. This is critical for the validity of syllable decoding results from MEG data.

Participants’ performance on the *overt* version of the task is summarized in **Table S1** (see also **Fig S1**). The median sound onset of syllable 1 (event 1) occurred 439 ms after the presentation of the syllable on the screen (*Methods*). The onset for syllable 2 (event 2) occurred on average 175 ms after the fixation cross. These times were taken to suggest the *expected imagery onset*, under the assumption of similar timing during the *Imagery* condition. Importantly, the interquartile range for the two events (syllable 1: 99 ms; syllable 2: 146 ms) indicated that participants were more precise in time in the production of the first imagined event than the second. We expected these differences in variability to have an impact on decoding, with a greater alignment for event 1 translating into better decoding.

Since, by definition, there is no overt behavioral output of imagery, a first step was to measure and quantify the difference between the *Imagery* condition and a control condition matched in all respects. In this *Reading* condition, participants simply passively looked at the syllables and fixation crosses appearing at the center of the screen. To evaluate the extent to which *Imagery* contained information over and above *Reading*, we used a decoding approach (temporal generalization; **Fig 2**). In addition to quantifying existing differences between the two conditions, this approach can track the dynamics of neural processes underlying a particular experimental condition^42,43^. Therefore, to track the development of neural processes common to speech imagery (i.e., processes shared by the three imagined syllables) as distinct from passive viewing/reading, we trained a linear classifier on the *Imagery* versus *Reading* contrast at each time point and tested its performance across all timepoints within the trial. This analysis was performed for each subject separately, using stratified 4-fold cross-validation with regularization and Receiver Operative Characteristic Area Under the Curve (ROC AUC) as the scoring metric (*Methods*). The analysis resulted in a temporal generalization (TG) matrix per subject, which we then averaged across subjects. We expected areas of higher decoding accuracy (i) at the expected imagery onsets (as determined from participants’ overt productions during training; **Fig S1**; **Fig 3A** upper panel; Table S1) and (ii) before imagined event 1, for motor planning of the syllable to be produced in each trial (**Fig 2**).

**Fig 2.**
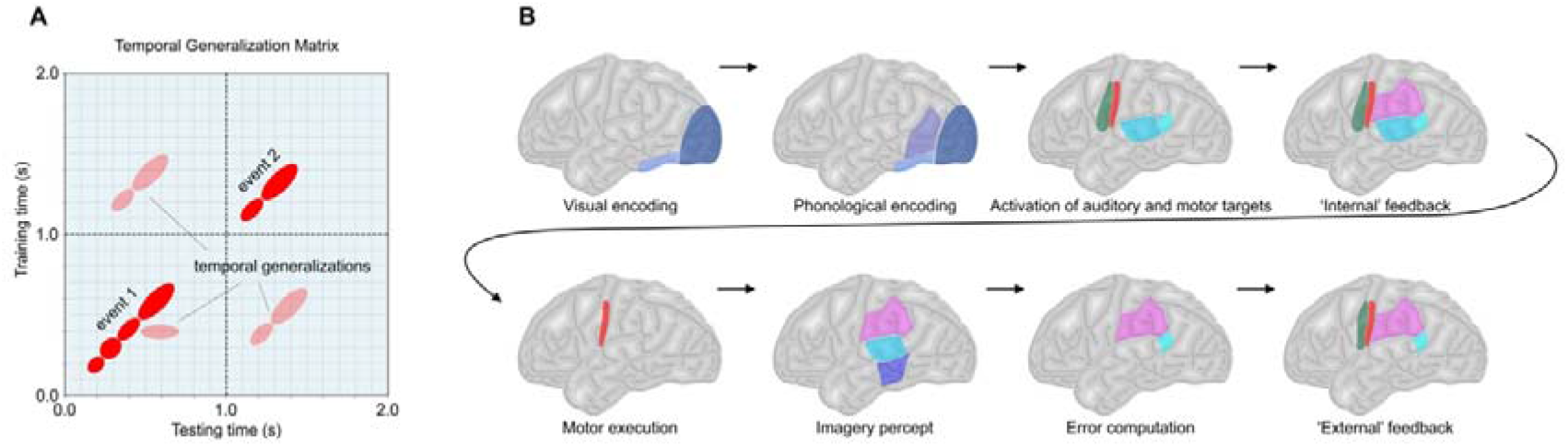
Main experimental hypotheses. **A. Temporal Generalization.** To discover and quantify syllable decodability in imagery, we used a multivariate pattern analysis in which classifiers are trained on a single time sample (rows; *y*-axis) but tested on all time samples of the trial (columns; *x*-axis). This results in a temporal generalization (TG) matrix that depicts the extent to which a given neural pattern is present across time. The TG method is a powerful approach to reveal not only the number and approximate times of neural processes but also the nature of the underlying representations (e.g., evolving, reactivated, ramping; see^42,43^ for a full explanation of the method). We expected high decoding accuracy (red circles/ellipses) if neural processes underlying speech imagery instantiate decodable representations. The schematic depicts four distinct processes for event 1 with some temporal generalization and two for event 2 as an example. High accuracy was expected for distinct neural processes underlying both imagery events (event 1 and event 2). We also expected event 1 representations to generalize to event 2 and vice versa. **B. Expected sequence of neural processes for speech imagery**. Insofar as speech imagery mirrors overt speech, we hypothesized that the underlying sequence of neural processes should conform to current models of speech production. Inspired by a recent model^8,9,45^ derived from other areas of motor control, we predicted 1. A *speech planning stage* encompassing visual encoding (visual and inferotemporal cortex), phonological encoding (left posterior middle and superior temporal cortex), the parallel activation of auditory and motor targets (auditory and motor cortex, respectively), and an ‘internal’ feedback loop for error correction characterized by sensory-motor activity; and 2. A production stage involving motor execution (motor cortex), activation of the imagery percept (auditory and somatosensory areas), prediction error elicited by the lack of overt auditory feedback (posterior auditory cortex); and a second feedback stage (‘external’ feedback) again with sensory-motor interactions. Note that since we are assessing *imagery*, rather than overt speech, the common nomenclature of *internal* and *external* loops, which refers to the nature of the feedback, is not strictly applicable. However, we decided to keep this nomenclature to link our predictions and results to theoretical formulations of the model.

**Fig 3.**
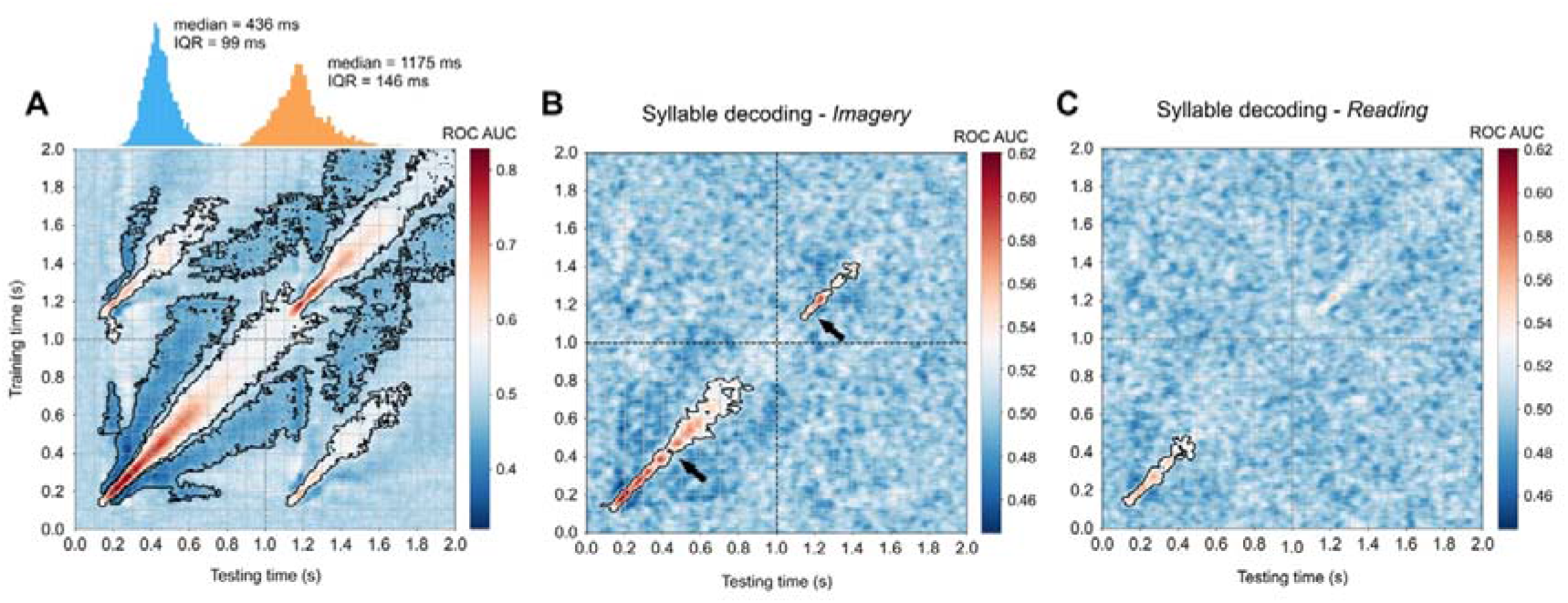
Temporal generalization matrices track the sequence of neural representations during speech *Imagery* and *Reading*. **A.** Average TG matrix (N = 21) for the contrast *Imagery* vs. *Reading* as well as syllable onset time distributions (top inset) for all syllables *overtly* produced during training by all participants (blue = event 1; orange = event 2), plotted for reference as to the expected imagery onset. IQR = interquartile range. ROC AUC = Receiver Operative Characteristic Area Under the Curve (chance = 0.5). **B**. and **C**. TG matrices for the pairwise contrasts between syllables (*pa* vs. *ka, pa* vs. *ta*, and *ta* vs. *ka)* for each condition (*Imagery* and *Reading*, respectively) first averaged within subject and then across subjects (N = 21). Black arrows indicate the median syllable onset time of participants’ overt productions during training (event 1 median: ~436 ms; event 2 median: ~1175 ms). Clusters of statistically significant decoding (*p* < 0.05; black contour lines) were in all cases determined at the second level of analysis via a cluster-based permutation test across subjects (1000 permutations; two-tailed). Statistical significance indicates consistency across subjects, while high ROC AUC values reflect robust classifier performance on discriminating the contrasts.

**Fig 3A** shows the average TG matrix across the sample. Clusters of statistically significant decoding (*p* < 0.05; black contour lines) were determined at the second level of analysis via a cluster-based permutation test (1000 permutations; two-tailed) across subjects. Large regions of high decoding accuracy (with ROC AUC values up to 0.82) suggest robust and consistent differences over time between *Imagery* and *Reading* conditions. The extended nature of the clusters suggests that the differences are driven both by domain-general processes (such as, perhaps, attention), as well as processes specific to speech imagery (i.e., bearing speech content). As expected, a significant degree of generalization is observed between the two imagery events within the trial, indicating similar neural underpinnings.

### Decoding participants’ imagined speech from MEG data

Having established that the MEG signals during *Imagery* contain information distinct from *Reading*, we next asked whether *Imagery* trials carried discriminable *content* regarding the three imagined syllables (/pa/, /ta/, and /ka/). Here, the TG approach can provide *direct* evidence for speech imagery, if areas of significant decoding are found, but it also provides valuable insight into the nature of the neural processes involved, for example, in terms of the number of distinct processes, their times of occurrence, and generalizability (**Fig 2**). We first generated, for each participant and condition, a TG matrix for each of the pairwise syllable contrasts (/pa/ vs. /ka/, /pa/ vs. /ta/, and /ta/ vs. /ka/) following the same decoding approach as before. These three matrices per participant and condition were then averaged within subject and entered into a cluster-based permutation test across subjects (N = 21; 1000 permutations; two-tailed) to determine clusters of significant syllable decodability (*p* < 0.05) in each of the conditions (*Methods*). Given that the processing of visual information (i.e., reading the syllables) is shared between *Imagery and Reading*, we expected this analysis to yield some similarities in the early stages of event 1. However, direct evidence for speech imagery would also require for syllable decodability to extend further in the *Imagery* condition only in event 1 and stand alone in event 2, reflecting the occurrence of the actual speech imagery events.

Clusters of relatively high decoding accuracy (ROC AUC scores up to 0.62, significant at *p* < 0.05) during event 1 reveal a distinct cascade of neural processes during *Imagery* (**Fig 3B**). The limited span of these successive clusters both on and off diagonal indicates that the representations involved were rapidly evolving (50 ms - 60 ms) and highly specific (with limited generalization). This result merits special emphasis: it is possible to decode with high time resolution a participant’s internally produced and minimally contrasting syllables from non-invasive data. Significant decoding starts immediately after syllable presentation (~120 ms) and extends well beyond the expected imagery onset (**Fig 3B** black arrow at ~436 ms). Syllable decodability during event 2 was sparser, albeit significant in clusters immediately before and after the expected imagery onset (**Fig 3B** black arrow at ~1175ms). This replicates the decoding results for event 1 and at times when imagery is only prompted by a fixation cross. The sparser decoding for this event and the absence of generalization between events likely reflect the temporal misalignments within and between participants’ inner productions, as suggested by their overt productions (**Fig 1**; **Fig 3A**). As predicted, syllable decodability in the *Reading* condition was significant, if weak, in clusters between ~120 ms and ~450 ms (**Fig 3C**). This suggests a similar succession of neural processes to the *Imagery* condition up to the imagined event, or extended visual processing during *Reading*. In favor of the latter interpretation, no clusters of significant decoding were found during event 2 in the *Reading* condition when no visual information of the syllable was present.

An important concern and classic critique of imagery studies derives from the associated, if small, movements that can accompany mental imagery. We took great care to control for that potential confound. The analysis of the EMG data by participant indicated, as expected, the presence of micromovements (**Fig S2**-**Fig S4**). Micromovements are a common phenomenon during imagery and inner speech and are commonly assumed to be a byproduct of motor signals that cannot be fully inhibited^7^. Interestingly, micromovements were present both in the *Imagery* and *Reading* conditions. We performed an in-depth analysis of the EMG data to ensure our decoding results could not be explained by participants’ micromovements. Although small differences were found between *Imagery* and *Reading* conditions (**Fig S2** and **Fig S4**), the micromovements did in no case discriminate between the imagined syllables (**Fig S2** and **Fig S4**).

### Neural dynamics underlying speech imagery

So far, we were able to decode participants’ speech imagery and uncover a series of well-defined stages leading to the imagined speech event. We next sought to establish the neural correlates of these stages. On the one hand, this analysis can adjudicate between theories of speech imagery that posit a close parallel between imagery and overt production and theories that conceptualize imagery as a byproduct of motor planning (i.e., without primary motor involvement)^49^. On the other hand, if imagery mirrors overt speech, the sequence of neural events underlying imagery can adjudicate between current models of speech planning and production (**Fig 2B**).

To characterize the neural dynamics underlying speech imagery, we first acquired structural MRI data from a random subsample of participants in the *pa-ta-ka* cohort (*Methods*). Each participant’s *Imagery* condition’s average time series was projected to their native source space and then morphed to a common coordinate space (Montreal Neurological Institute) before averaging across participants (*Methods*). The goal of this group analysis was to assess the sequence of neural activity that gives rise to speech imagery.

**Figure 4** shows the progression of neural activity during *Imagery* between 120 ms and 610 ms after syllable presentation, corresponding to the islands of significant syllable decoding previously identified. The analysis was thus guided by the syllable decoding results (**Fig 3B**) since these inform us about the times of relevant mental representations within the trial. (See **Fig S6** for the complete sequence of neural events.)

**Fig 4.**
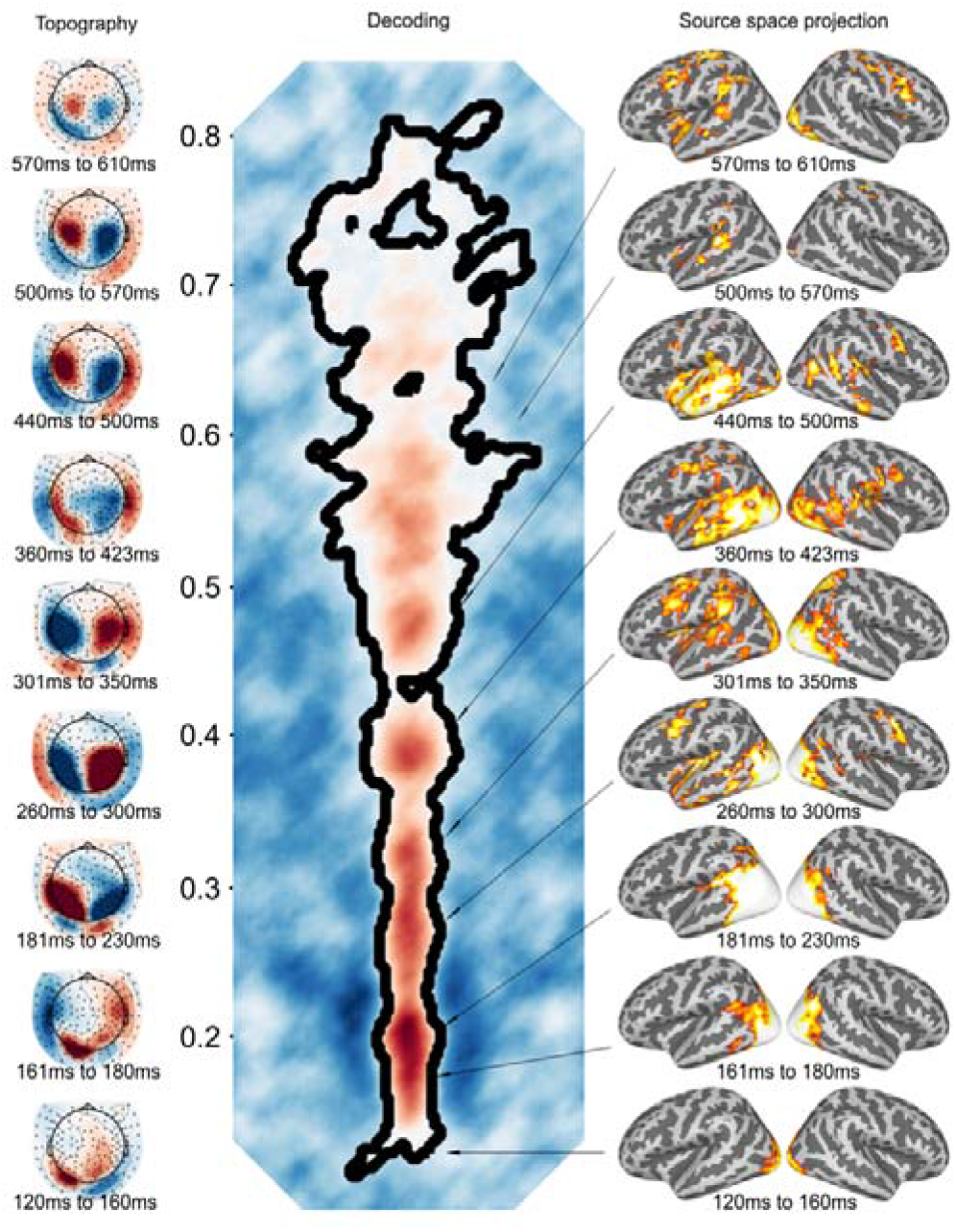
Clusters of syllable decodability reveal a processing cascade during speech *Imagery* consistent with SFC models. Response topographies (*Imagery*) averaged over participants (left panel) and activity for a subgroup of participants estimated with sLORETA (right panel) corresponding to the clusters of significant syllable decoding in the *Imagery* condition, event 1 (center panel; from Fig 3). Response topographies were thresholded at +-20fT. Source space activity was thresholded at minima ranging between 1.88 and 3.34 and maxima between 2.37 and 4.51 units for display purposes. Time progress from bottom to top.

Both source and sensor renderings show a clear sequence of distinct neural events, starting with visual areas (~120 ms) for visual encoding. This activity extends to the well-established ventral visual pathway and subsequently to the lateral temporal cortex (~180 ms), particularly in the left hemisphere, putatively reflecting a phonological (re)coding of the syllables from their visual form. This stage is followed by activity in the auditory cortex, the Sylvian parieto-temporal area, frontal anterior insula, and bilateral pre-motor regions between 260 ms and 300 ms, which may relate to auditory-motor integration processes and speech motor planning^9,47,50,51^. Extensive activity over (predominantly left) auditory regions can be seen after 440 ms, that is, at and following the expected imagery onset (estimated at 436 ms for the entire sample and at 444 ms for the MRI group; Table S1). We conjecture this activity to be the neural correlate of the quasi-perceptual experience that defines speech imagery, in line with previous research^11,15,18^. The decoding clusters indicate that this imagery event is flanked by two additional distinct stages. First, a stage prior to (internal) production (~300 ms to ~400 ms) featuring activity in both premotor/motor and temporo-parietal regions. This is consistent with the internal feedback loop of SFC theories, characterized by feedforward-feedback processes for speech planning^8,9,45^; and second, a stage following production (>500 ms) featuring activity in posterior auditory as well as bilateral motor regions, consistent with a hypothesized feedback stage following motor execution^8,9,45^. As might be expected, the same analysis performed subtracting the *Reading* condition from the *Imagery* condition in source space removes much of the visual activity, but the exact same sequence can be observed (**Fig S7**). Note that this is a very conservative analysis controlling for neural processes not specific to imagery. The result of this analysis also provides additional evidence that syllable decoding in the *Reading* condition was based on the visual encoding of the syllables (visual representations) throughout, in contrast to the *Imagery* condition. Event 2 did not yield as clear a sequence (**Fig S8** and **Fig S9**), as expected from the misalignments within and across subjects and the sparser decoding results. It is nevertheless worth highlighting the implication of auditory, premotor, and motor areas as well as of the Sylvian parieto-temporal area in this latter event, suggesting similar neural dynamics to event 1.

### The internal and external loops in State Feedback Control

Our analyses revealed a sequence of neural representations leading to a speech imagery event consistent with the SFC architecture (**Fig 2**). A critical feature that distinguishes SFC from other models of speech production is the existence of both an internal and an external feedback loop supported by the interplay between sensory and motor regions planning^8,9^ (**Fig 5A**). The internal feedback loop, by hypothesis, is critical for speech planning in its ability to issue corrections, *before motor execution*, for discrepancies between the predicted sensory outcomes of planned actions (i.e., somatosensory and auditory predictions) and the intended actions/sounds (i.e., somatosensory and auditory targets). Note that this comparison is contingent on the early activation of the sensory targets, a feature that is also absent in the DIVA architecture. The external loop, in contrast, issues corrections for discrepancies between predicted and *actual* sensory outcomes (i.e., post execution) which can then be used to fine-tune the internal forward model used to generate the sensory predictions.

**Fig 5.**
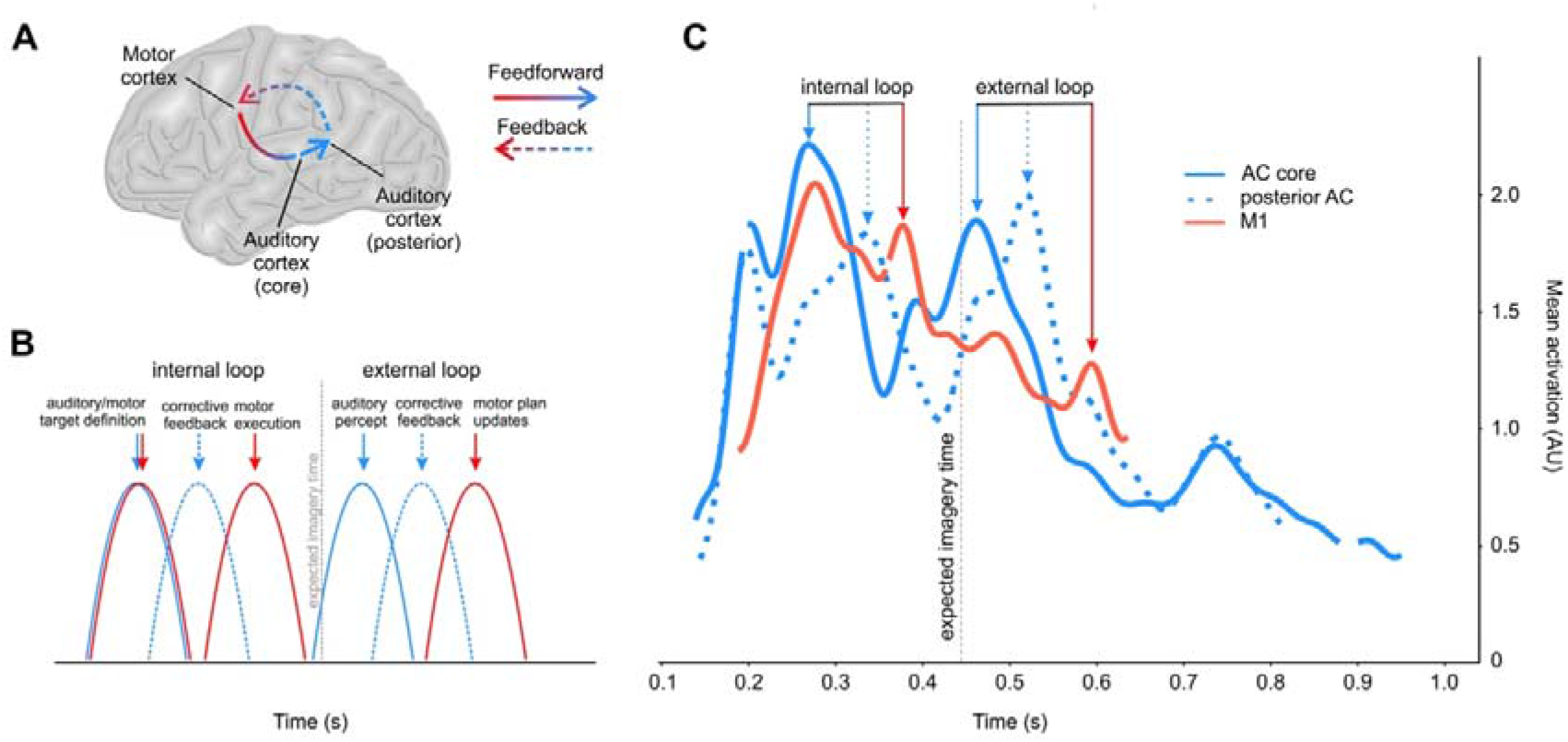
Time courses of auditory and motor regions during speech imagery. **A. Feedforward feedback loop for speech motor control.** SFC theories^8,9,45^ hypothesize an internal loop for motor planning and an external loop for post-production feedback, both characterized by feedforward and feedback processes between motor and auditory regions. Predictions from speech motor plans about their auditory consequences are compared with auditory targets (feedforward arrow). Corrective error feedback is sent back to motor regions to refine motor plans (feedback arrow). **B. Schematic of expected peaks of activity in auditory and motor regions.** SFC posits the parallel activation of auditory and corresponding motor plans (auditory and motor targets). A forward prediction of the consequences of the selected motor plans is compared to the auditory target, generating an error signal commensurate with the discrepancy. The error signal is sent back to and used as corrective feedback by motor regions, which ‘execute’ the motor command. Internal motor execution immediately preceding the expected imagery onset generates activity in auditory regions (a second feedforward sensory prediction) likely associated with the auditory percept (imagery). Since there is no feedback from overt production to match this sensory prediction, a second error signal is generated and fed back to motor regions. **C. Time courses of auditory and motor regions.** The time courses of core auditory and motor regions in addition to a posterior auditory region known for its role in sensory feedback show activity pre and post internal speech production (vertical dashed line) consistent with the hypothesized double feedback loop. Blank segments in the lines indicate times of non-significance. Time courses were low-pass filtered at 20 Hz (double pass Butterworth of order 5) for display purposes.

Although the previous analysis clearly shows activity in motor and sensory regions, both prior to and following the expected time of inner production, that analysis represents only a snapshot of their dynamics. To examine these dynamics more closely, we extracted the time courses spanning event 1 from three key regions of interest (ROIs) in the left hemisphere, namely motor cortex, primary auditory cortex (core), and posterior superior temporal auditory cortex (posterior auditory) (*Methods*). Besides their hypothesized role in the internal feedback loop, this restricted ROI selection was motivated by the widely agreed-upon functions of these regions^52^: it is reasonable to attribute motor representations to motor cortical areas and auditory representations to auditory areas. Note that, in addition to a core auditory region, we selected a posterior auditory region for its known involvement in the computation of auditory feedback^53^. The anatomical location of the 3 ROIs was based on a well-known cortical atlas (Glasser et al.^54^) (see *Methods* for the detailed procedure of the selection of the ROIs). In short, we selected, for each participant, the MNI coordinates that displayed maximal activity within each of the corresponding atlas labels. Around each coordinate point, we then built a 4 mm sphere and extracted the average time course of the sources within that local volume. We determined the times at which time courses therein were consistently activated across participants (significantly above their mean baseline activity; *Methods*) using a cluster-based permutation test (1000 permutations; one-tailed). We expected the sequential activation of core auditory (coding for the acoustic representation), posterior auditory (coding for the prediction error), and motor areas (receiving feedback) to occur *at least once* before the expected time of imagery (i.e., during speech planning), with a similar pattern also occurring after internal production (**Fig 5B**).

**Fig 5C** shows the time courses of the selected ROIs (see **Fig S10** for the activity of control regions, **Fig S11** for additional auditory regions, and **Fig S12** for somatosensory regions).

The time courses show an initial peak of activity in auditory areas (~180 ms), in line with an initial phonological (re)coding stage, followed by concurrent activity in both motor and core AC regions. This activity is consistent with the parallel activation of motor and auditory targets hypothesized by SFC models. This is followed by activity in posterior auditory regions and motor activation immediately before the expected imagery onset (vertical dashed line), consistent with an inner feedback stage and internal motor execution. Following motor activity, core auditory activity rises to peak at the expected time onset. We conjecture that this auditory activity (plus activity in secondary auditory regions; **Fig S11**) reflects the inner ‘hearing’ of the imagined spoken syllable. The time courses also show that core auditory activity at the expected imagery onset is closely followed by posterior auditory and motor activity, consistent with the second feedback loop hypothesized by SFC. In all, the dynamics of auditory and motor regions are consistent with an internal feedback loop prior to motor execution as well as an external feedback loop following motor execution, both attributes predicted by SFC theories^8,16,44^. The time courses of control ROIs (e.g., left visual cortex and frontal pole; **Fig S10**) show that these auditory and motor group-level dynamics are not a product of the analytical procedure.

### The nature of the representations in speech imagery

To validate the temporal dynamics of speech imagery and to extend the results beyond the phoneme contrasts we employed (/*p*/, /*t*/, /*k*/), we replicated the experiment using a new participant cohort (N = 9; *Methods*). This time, however, we asked participants to imagine a different set of syllables, specifically /ta/, /tu/, and /ti/, varying in their vowels rather than in the consonants, and therefore with a higher contrast in the acoustic than in the articulatory domain. Accordingly, we should see that the decoding cluster immediately preceding the expected imagery onset, which corresponds to motor/somatosensory activation (**Fig 4**) and thus presumably reflects the contrast between motor or somatosensory representations, is relatively de-emphasized (i.e., less discriminative of the syllables’ identities) in the *ta-tu-ti* set, since the initial position of the articulators for the three syllables should now be more similar (**Fig 6A** top). In contrast, we should find that the decoding cluster immediately following the expected imagery onset, which corresponds to auditory activity and presumably reflects the contrast between auditory representations, is now relatively emphasized (is more discriminative of the syllables’ identities) and extended, echoing the more prolonged nature of the vowel contrast in auditory space (**Fig 6A** bottom). **Fig 6B** shows the results of the syllable decoding analysis for this new set of syllables. (See **Fig S13** for the *Imagery* vs. *Reading* contrast and the syllable decoding analysis in the *Reading* condition).

**Fig 6.**
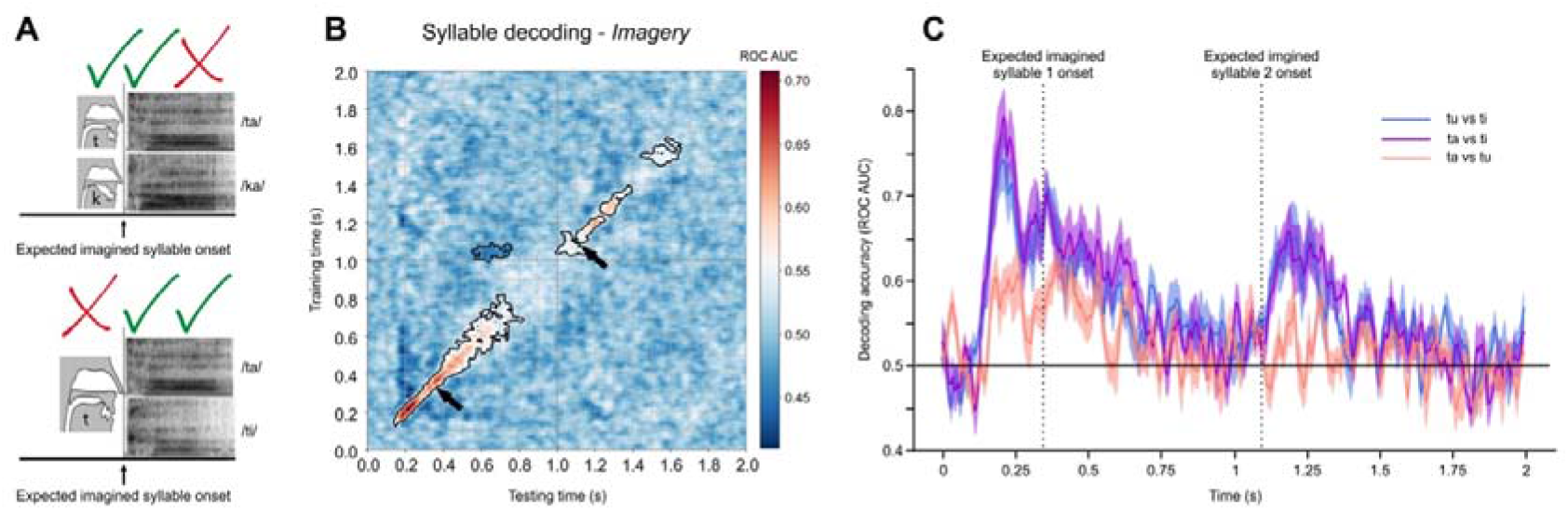
Analysis of the *ta-tu-ti* set. **A**. Schematic of the predictions for the syllable decoding for the *ta-tu-ti* set compared to the *pa-ta-ka* set. Top: The main decodeable contrasts for syllables /pa/, /ta/, and /ka/ are the initial position of the articulators (motor and/or somatosensory representations) and the consonant-vowel transitions (motor, somatosensory, or auditory representations) (Schematic shows /ta/ vs. /ka/ only). Bottom: In contrast, the main decodeable contrasts for syllables /ta/, /tu/, and /ti/ are the consonant-vowel transitions (motor, somatosensory, or auditory representations) as well as the stable parts of the vowels (Schematic shows /ta/ vs. /ti/ only). **B.** TG syllable decoding matrix (averaged across contrast and subjects). Black arrows indicate the median syllable onset time of participants’ overt productions during training (event 1 median: ~349 ms; event 2 median: ~1098 ms). Clusters of statistically significant decoding (*p* < 0.05; black contour lines) were in all cases determined at the second level of analysis via a cluster-based permutation test across subjects (1000 permutations; two-tailed). **C.** Pairwise syllable decoding contrasts (/tu/ vs. /ti/, /ta/ vs. /ti/, and /ta/ vs. /tu/). Shaded regions show the standard error of the mean. Vertical dashed lines indicate the expected imagery onsets for each of the imagery events. Solid black line indicates chance level.

Several observations can be drawn from these new decoding analyses (**Fig 6B** and **Fig S13**). First, these new results validate our previous decoding results (for *pa-ta-ka*) and indicate that the approach generalizes beyond the original syllables employed. Second, as predicted, the decoding in the *ta-tu-ti* set prior to the expected imagery onset was relatively weaker than the decoding following the expected imagery onset. That is, higher accuracies were obtained at the times when we expected the vowel sound contrast (**Fig 6BC**) than at the times corresponding to the consonant articulation, since this was common across the syllables. This is in line with the hypothesis that motor and auditory representations are what are primarily identified by the decoding analysis of *pa-ta-ka* and *ta-tu-ti*, respectively, and corresponding to the times immediately before and after the expected imagery onset. The fact that the pairwise contrasts with the syllable /ti/ were the more robust (**Fig 6C**) represents additional evidence for the idea that the decoding in the *ta-tu-ti* set primarily leverages acoustic representations, since the distances in acoustic space between /ta/ and /ti/ and between /tu/ and /ti/ are greater than the distance between /ta/ and /tu/^55^. (See **Fig S14** for the pairwise contrasts in the *pa-ta-ka* set). Third, although the contrasts between *Imagery* and *Reading* were comparable between the two syllable sets, the imagined syllables involving vowel contrasts (/ta/, /tu/, and /ti/) were overall more accurately decoded from the neural data than syllables involving consonant contrasts (/pa/, /ta/, and /ka/) (**Fig S15**). This indicates that, although the decoding of motor representations pertaining to imagined syllables is possible from non-invasive data, the decoding of auditory representations may be more robust, perhaps due to their more prolonged rather than transient nature, which may partly correct for trial-to-trial misalignments. And fourth, the identified motor/somatosensory and auditory representations in both syllable sets collectively span at least a full syllable’s length (>200ms). This contrasts with the rapidly evolving (compressed) representations during speech planning (in the order of 50 – 60 ms; **Fig 3** and **Fig 4**).

## Discussion

Speech imagery is the capacity to ‘hear’ self-generated (covert, internal) speech. Despite its pervasive nature in many aspects of cognition, the study and use of speech imagery in both clinical and academic research has been challenging. We characterized the dynamics of a relatively simple speech imagery task, by pairing neurophysiological (MEG) data with a novel experimental protocol designed to overcome the known methodological difficulties. Using a time-resolved decoding analysis of participants’ imagined utterances, we show that generating imagined speech involves the rapid succession of relatively encapsulated neural representations that map in systematic ways onto some predictions of current theoretical models of overt production. In particular, the data support the state feedback control conceptualization of speech production.

The neural correlates we report for these transient representations are consistent with previous fMRI research on speech imagery, inner, and covert speech^7,10,56^. An advantage of the time-resolved approach we pursue is that it provides, in addition, a dynamic characterization of speech imagery, which brings us closer to mechanistic understanding. Broadly, the data are consistent with two well-defined stages during imagery, namely planning and (internal) production (**Fig 4**). The production stage is characterized by widespread left-lateralized auditory activity at the expected syllable onset (~400 ms, based on participants’ overt productions of the same syllables), immediately preceded by speech motor activity. We hypothesize that this auditory activity corresponds to the percept associated with speech imagery^11,18^. Following the imagined event, the syllable decoding approach identified two additional time periods with distinct neural representations, associated with pSTG activity (~500 ms) and subsequent bilateral pre/motor activity (~570 ms). This pattern of activity is consistent with a hypothesized external feedback loop during overt production, in which errors from comparing the predicted sensory consequences of planned articulatory gestures with their actual consequences (i.e., auditory feedback) are forwarded to motor regions^8,45,47^. Indeed, the location of the posterior auditory cluster is consistent with fMRI research using altered feedback to identify error-related activity^57,58^ as well as recent ECoG data distinguishing sensory processing (in more anterior regions) from feedback error signals^53^. Activity in bilateral pre/motor regions has also been reported following auditory errors^57,59^, in line with our observations. Interestingly, while error-related activity can be modulated experimentally by altering feedback in both overt^57,59^ and imagined speech^13^, it is unclear what the expectation should be in the case of imagery when no matching or mismatching inputs are given. In accord with a recent hypothesis on predictive processing^20^, our intuition is that, since there is no overt auditory feedback to meet predictions (i.e., there is less input than predicted), an error response in auditory regions should still be produced. Our results support this hypothesis, highlighting the potential of our approach for research on predictive processing.

Regarding speech planning, we identified at least three distinct time periods of significant syllable decoding prior to the production stage (**Fig 4**). The first, ~180 ms after syllable presentation, was associated with activity in left posterior temporal cortex and thus consistent, both in time and location, with a much-theorized phonological encoding stage^9,50,60^. Phonological encoding may thus be present whether production is internally generated (e.g., from abstract thought) or externally triggered (e.g., from reading). The second was characterized by concurrent activity in left auditory and bilateral motor regions at ~260 ms. Such activity is predicted by SFC models recently applied to speech motor control in which auditory and motor targets are accessed in parallel immediately following phonological encoding^9^. SFC also predicts an internal feedback loop prior to motor execution, in which sensory predictions (auditory and somatosensory) are compared to the intended sensory targets^8,9,45^. Although theoretically well-grounded, there is no direct empirical support for an internal loop in speech production. Indirect evidence comes from the timing with which individuals correct themselves during speech errors, which are too fast for responses to external auditory feedback^61^. Activity at ~300 ms (**Fig 4)** is consistent with the hypothesized internal feedback loop, featuring the posterior auditory cluster, the supramarginal gyrus, and pre/motor regions. This result is further supported when examining the time courses of auditory and motor areas during imagery (**Fig 5**). Specifically, the sequence of activations observed for the putative external loop is also present immediately before the expected time of imagery, consistent with the monitoring and planning role of the internal loop. A tentative hypothesis is that the observed activity in ventral somatosensory and supramarginal regions (**Fig S12**), which closely mimics that of core and posterior auditory areas (respectively), reflects a level of speech motor control additional to that subserved by auditory and motor interactions, in line with hierarchical SFC models^9^. By this account, inferior parietal activity prior to speech imagery events previously reported in the literature (e.g., ^11,15^) could relate to this level of control.

The generation of speech imagery appears to closely mirror that of overt speech, with full-blown planning and (internal) execution stages. Our data are thus inconsistent with views of imagery as a by-product of motor planning (e.g., ^49^). Moreover, given the presence of unspecific micromovements at the expected imagery onsets (**Fig S2-S4**), our hypothesis is that, during imagery, speech plans are executed but aborted (inhibited) at the periphery. In this sense, speech imagery may be seen as analogous to concrete (as opposed to abstract) forms of inner speech^7^.

The properties of the neuroimaging method (MEG) invite care with regard to the accuracy of the reported cortical areas. However, although possible in principle, we suspect major inaccuracies are unlikely given the close correspondence between the sources of activity in our imagery task and previously reported clusters in motor and sensory areas^56^. This is particularly so in the case of auditory regions, which show a consistent pattern of activity at the expected time (**Fig S10**). A possible exception could be the posterior auditory cluster because of its proximity to area Spt (cf., ^62^). Indeed, the behavior of this cluster could be seen as consistent with Spt’s hypothesized role in auditory-motor transformations^8,63^. We also found activity in premotor and motor regions to be in very close spatial proximity (**Fig 4**). It is therefore possible that the activity we attribute to the motor cluster pertains to premotor cortex instead. This would be consistent with a hypothesized origin of sensory predictions in the *premotor* cortex in response to inputs (efference copies) from motor areas as well as with known inputs of premotor areas to motor cortical regions for production^47^.

A further noteworthy finding revealed by the decoding analysis is the *compressed* nature of speech representations during planning. These representations appear to change rapidly, every 50 to 60 ms, which contrasts with the natural rate at which the internal productions unfold (cf., ^64^). There is evidence that inner speech and speech imagery percepts encode tempo, pitch, timbre, and loudness information^13,14,39,65^ but little is known about the relationship between production and planning stages in imagined, inner, or overt speech. This result is particularly relevant for potential brain-computer interfaces, which could account for this feature to increase decoding performance.

Finally, we highlight some methodological considerations. In decoding imagined speech, we would like to emphasize the importance of aligning responses to reduce the amount of noise in the data with which classifiers are trained. We took several measures to improve the alignment of imagined events, both within and between participants (*Methods*). Many difficulties in decoding imagery from broadband signals are due to temporal misalignments. Indeed, many speech decoding studies using time-resolved methods (e.g., EEG, MEG) have ultimately resorted to frequency analyses (e.g., power modulations, cross-frequency coupling)^66^, loosing temporal resolution. While our objective was not decoding per se, we were able to decode participants’ imagined utterances with relatively high accuracies from non-invasive data using simple linear classifiers. Although we are optimistic about recent invasive approaches and sophisticated analysis pipelines (e.g., ^41^), we would like to emphasize that non-invasive methods and ‘lighter’ analytical procedures can also provide systematic insight.

In sum, combining traditional non-invasive methods and state-of-the-art analytical techniques, we show how speech imagery works. This approach opens a wide range of possibilities both for basic and clinical research. As an example, we show that speech imagery closely mirrors overt speech planning and production with an underlying sequence of neural events consistent with current models of speech motor control, and involving (minimally) both motor and auditory representations. Additionally, we expose the contrast between the compressed nature of these representations during planning and the natural speed at which they unfold during the internal perception of the imagined speech event. These findings also sound a note of hope for the development non-invasive speech interfaces.

## Methods

### Participants

A total of thirty subjects participated in the study. Twenty-one (15 women; mean age = 28.19 years; *std* = 6.57) were tested on syllables *pa*-*ta*-*ka*. From this cohort, we acquired structural MRI data from a random subsample of 10 participants (5 female, mean age = 31.8, std = 6.26). An additional 9 participants (7 women; mean age = 23; *std* = 7.94) were tested on the syllables *ta*-*tu*-*ti*. All participants were right-handed and self-reported no neurological deficits. They also provided written informed consent before participation. The protocol was approved by the local institutional review board (New York University’s Committee on Activities Involving Human Subjects).

### Task

The speech imagery task (Fig 1) involved imagining (internally producing) a different syllable on every trial. Two sets of three different syllables (*pa*, *ta*, or *ka*; and *ta*, *tu*, *ti*) were tested, each on a different cohort. The task was identical otherwise. Each trial began with a fixation cross of variable duration (1-1.5 sec) in the center of the screen. One of three syllables tested (*pa*, *ta*, or *ka*; or *ta*, *tu*, *ti*) was presented and remained on the screen for exactly 1 second. Syllable presentation was followed by another fixation cross lasting 2.5 seconds, at which point the next trial began. The experiment comprised two conditions, *Imagery* and *Reading*, each with 4 blocks of 120 trials (40 presentations of each syllable per block), counterbalanced between participants. The total number of trials over the course of the experiment was 960 (320 per syllable). Syllable presentation was fully randomized within each block. In the *Imagery* blocks, participants were instructed to imagine producing the given syllable as soon as possible after its appearance on the screen (event 1) and a second time on the fixation cross (event 2). Each trial, therefore, comprised two imagined events (event 1 and event 2). In the *Reading* blocks, participants were instructed to passively look at the center of the screen. The task was scripted and delivered using custom python scripts and Psychopy software.

### Pre-screening and training

Participants were required to take part in an informative and training session prior to the MEG session (1-7 days), in which the researcher/s explained the nature of the experiment and the desired type of imagery (i.e., production-based rather than hearing-based imagery). Participants were familiarized with the actual imagery task (Fig 1) and encouraged to practice briefly following the example of the researcher and using overt productions rather than imagery to be able to evaluate their grasp of the task. At this point only, participants were given feedback regarding the timings of each production (i.e., their timing precision). We finally asked participants to complete a pre-screening practice block in an isolated sound booth, also using overt production. Their productions were recorded for informal analysis. After the practice block, we plotted the onsets of their syllable productions and evaluated the adequacy of the participant for further testing based on the dispersion of their distributions. To ensure a minimum consensus across the cohort, participants with large IQR (>200ms) in either imagined event were discarded for further testing. Participants with smaller IQRs were invited to participate in the MEG session. We additionally provided participants with an online version of the task and encouraged them to practice in their own time. Lastly, each participant additionally completed a minimum of 1 practice block on the day of the MEG acquisition using overt productions which were recorded for subsequent analysis (see below).

### Analysis of overt speech productions

Participants completed at least one overt practice block immediately before the MEG session in which they were instructed to speak the syllables aloud. In an isolated sound booth, participants sat in front of a computer with a microphone close to their mouth and performed the task while their productions were recorded. This produced an average of 120 wav files per participant which were subsequently analyzed using Praat. Each utterance (2 events per trial) was manually annotated for syllable and formant transition onsets and offsets by a research assistant blind to the purpose of the experiment. Durations for syllables and formant transitions were computed by subtracting offset times from onset times. We computed medians and interquartile ranges for onsets and durations for both syllables and formant transitions.

Syllables onsets were determined based on the characteristic spectrogram signatures of the noise burst for each of the unvoiced stop consonants used (i.e., *p*, *t*, and *k*). Unvoiced stop consonants (e.g., *p*, *t*, *k*) involve obstructions by the articulators in the vocal tract which have specific acoustic signatures corresponding to a silent period (closure) and a noise burst (release). In particular, unvoiced stop consonants show clear differences in the shape of the noise burst: /p/ shows a short-lived wide range burst (across all spectrogram frequencies) that has lower intensity; /t/ shows a more prolonged burst in the upper part of the spectrogram (above 4 kHz) that has higher intensity; /k/ shows an even longer burst in the lower part of the spectrogram (below 4 kHz) that also has higher intensity (Kent, Kent, & Read, 2002). Syllable offsets were determined based on the speech envelope of each syllable.

Formant transition onsets were determined based on the characteristic spectrogram signatures of the noise burst for each of the unvoiced stop consonants used (i.e., *p*, *t*, and *k*), as above. Transitions offsets were determined based on the characteristic deflection patterns in the second (F2) and third (F3) formants. Formant transitions provide important perceptual cues for the manner (F1) and the place (F2 & F3) of articulation. /pa/, /ta/ and /ka/ are all plosives so manner of articulation remains constant. In contrast, /pa/ is bilabial, /ta/ is alveolar and /ka/ is velar, which means that the consonants differ in terms of place of articulation. As for the place of articulation, therefore, the first formant (F1) always raises after a stop consonant for all places of articulation, while the second (F2) and the third (F3) formant shape varies according to the place of articulation. For /p/ there is a raise in both F1 and F2; for /t/ F1 remains constant while there is a fall in F2; and for /k/ there is a fall in F1 and a raise in F2. The exact shape of the formant transitions will vary depending on the neighboring vowel, which in our case was also kept constant (/a/).

### MEG data acquisition and preprocessing

We acquired neuromagnetic responses from participants using a 157-channel whole-head axial gradiometer (KIT, Kanazawa Institute of Technology, Japan) situated in a magnetically shielded room, with a sampling rate of 1000Hz. To monitor head position during the recordings, five electromagnetic coils were attached to the subject’s head. We registered the location of these coils with respect to the MEG sensors before and after each block of the experiment. Participants’ head shape was digitized immediately before the MEG recordings using a Polhemus digitizer and 3D digitizer software (Source Signal imaging) along with 5 fiducial points, to align the position of the coils with participants’ head shape, and 3 anatomical landmarks (nasion, and bilateral tragus), to further allow for the co-registration of participant’s MEG data with their anatomical MRI scans. An online band-pass filter (1Hz-200Hz) was applied to all MEG recordings.

Data preprocessing was conducted using custom Python scripts and MNE-python software^67^. Bad channels were first selected and interpolated using spherical spline interpolation. A least-squares projection was then fitted to the data from a 2-minute empty room recording acquired at the beginning of each MEG session and the corresponding component was removed. MEG signals were next digitally low-pass filtered at 40Hz using MNE-python’s default parameters with *firwin* design and finally epoched between −1000ms and 2000ms relative to the onset of syllable presentation. Detrending and baseline correction (−1000ms to 0) were applied to the epochs. Cardiac and ocular artifacts were also corrected via independent component analysis. The epochs resulting from these steps were visually inspected and remaining artifactual trials were discarded from further analysis.

The same procedure was also repeated for each condition separately. A 40ms moving average was applied to the MEG data after filtering for the temporal generalization analysis by conditions only (Fig. 3), as means to increase the signal-to-noise ratio given the smaller number of trials for each syllable. This was achieved by convolving the data with a flat kernel of such length.

### Temporal Generalization

To characterize the dynamics of neural activity during speech *Imagery* in relation to *Reading*, we used a TG approach. The analysis was performed at the single subject level (i.e., within-subject). Specifically, we trained a linear classifier to discriminate between *Imagery* vs *Reading* trials at each time point of the trial and tested its performance across all time-points (i.e., from the presentation of the syllable to 2000ms later). We used 4-fold stratified cross-validation to balance the proportion of each class in each fold, alpha = 1 regularization, and ROC AUC as a scoring metric. The result of this procedure was a TG matrix per subject showing each classifier’s performance over time. Individual TG matrices were averaged across subjects to produce a group-level TG (e.g., Fig 3a).

To characterize the dynamics of neural activity during speech *Imagery* and *Reading* separately (e.g., Fig 3b and 3c), we generated TG matrices for each of the pairwise contrasts between the tested syllables (e.g., *pa* vs. *ka*, *pa* vs. *ta*, and *ta* vs. *ka*) following the same procedure described above. The three matrices per participant and condition were then averaged within subject and the resulting TG matrices (one per participant) were finally averaged across participants. This resulted in a single matrix per condition (e.g., Fig 3b and 3c), characterizing the sequence of neural processes for each of the two conditions (*Imagery* and *Reading*).

Clusters of statistically significant decoding (p < 0.05) across participants were in all cases (Fig. 3abc) determined via a cluster-based permutation test (1000 permutations; two-tailed). Note that significant decoding indicates consistency across participants over time, while higher ROC AUC scores reflect higher decoding accuracy or discriminability between conditions.

### Structural MRI data acquisition and source reconstruction

A high resolution T1 MPRAGE image was acquired from a random subsample of the original cohort (N = 10) with the following scanning parameters: TR = 2400 ms; TE = 2.24 ms; flip angle = 8°; voxel size = 0.80 × 0.80 × 0.80 mm^3^; 256 sagittal slices; and acquisition matrix = 320 × 300.

Each participant’s *Imagery* and *Reading* condition’s MEG data were projected to their native source space. We used a forward model based on a 1-layer boundary element model and a minimum-norm inverse model (signal-to-noise ratio = 2; loose dipole fitting = 0.2; normal orientation) using a noise covariance matrix computed from all sensors averaged over a pre-stimulus baseline period of 1 second across trials. This inverse model per condition was applied to the participant’s corresponding evoked responses using standardized low-resolution brain electromagnetic tomography (sLORETA). Each participant’s source reconstructed data was then morphed to a common coordinate space (Montreal Neurological Institute) and the 10 participants’ data were finally averaged to produce a single time course per vertex in source space (5124 vertices in total) per condition.

Note that close similarity between the larger cohort’s topographies and both the sensor space topographies of the MRI sample and the classifier patterns (coefficients) for syllable decoding (cosine similarity tests, **Fig S5**) indicates that the source reconstruction for the subsample is directly relevant to imagery.

### Time courses of auditory and motor regions of interest

To visualize the interplay between motor and auditory regions during speech *Imagery*, we plotted their time courses over event 1. To arrive at these time courses, we performed the following steps. First, we used a well-known anatomical atlas^54^ to select the MNI sources pertaining to the left primary and secondary auditory cortex (A1; lateral and posterior parabelt), the superior temporal gyrus/sulcus (A4, A5, dorsal anterior and dorsal posterior superior temporal sulcus), the ventral premotor cortex (ventral and rostral BA6, 55b, and PEF), and the motor cortex (BA4), as determined by the labels of Glasser et al.^54^. Since the atlas parcellation is relatively coarse (e.g., the motor cortex is characterized as a single region, irrespective of effectors), instead of taking all sources within a label, we selected the MNI coordinate points that displayed maximal activity within each area’s time course normalized by the mean activity over time across the entire brain (i.e., the maxima relative to average brain activity at each time point). Around each coordinate point, we built a 4mm sphere and extracted the average time course of the sources within from the raw signal (i.e., non-normalized over time) of each participant (1 second long from syllable presentation to fixation cross, i.e., event 1). In cases where the foci of activity were less than 1 mm apart, these regions of interest (ROIs) were merged by intersecting their vertices to extract a single time course. These steps resulted in time courses for 6 ROIs (AC core, mid STG/STS, dorsal posterior STS, posterior AC, ventral premotor cortex, and motor cortex) per participant. We determined the times at which these ROIs’ time courses were significantly above their mean baseline (1 sec) activity using a cluster-based permutation test (1000 permutations; one-tail) across participants.

### EMG data acquisition and analysis

Participants were instructed to avoid any articulatory movements (jaw, lips, or tongue) during the MEG recordings. Muscle movements were continuously monitored using electromyography (EMG). We recorded the EMG signal from two electrodes. One electrode was placed at the intersection between the right zygomatic bone and the mandible, to record jaw movements. The other electrode was placed between the right lower lip and the chin, to record lips movements. The complete setup also included a reference electrode placed on the right mastoid bone and a ground electrode placed on the right wrist bone. Electrodes were connected to an MEG-compatible BrainAmp DC amplifier (Brain Products GmbH, Gilching, Germany). The EMG signal was recorded at a sample rate of 500◻Hz. A 60 Hz notch filter was applied online to remove power line noise. Data was referenced online to the right mastoid. Electrode impedance was kept below 25◻kΩ.

EMG signals were filtered using a zero-phase, two-pass Butterworth bandpass filter with a 1 Hz high-pass frequency cut-off and a 50 Hz lowpass frequency cut-off. We segmented the raw signal into epochs between −1s before syllable onset and 2s after the syllable cue onset. Time series were down sampled to 250 Hz and the EMG signal was detrended. A baseline correction was applied by computing the mean of the 1s baseline period preceding syllable cue onset and subtracting this mean from the entire trial epoch. We used an auto-reject algorithm (Jas et al., 2017) for automatic artifact rejection. This algorithm defines a global threshold for artifact rejection that is specific for each participant based on a cross-validation procedure. This choice was motivated by the variability across subjects in the EMG data introduced by differences in skin conductance, muscle artifacts, heartbeat artifacts, prominence of the bones, etc… On average, 393.03 (± 105.75 std) *Imagery* trials and 390.48 (± 90.72 std) *Reading* trials survived the artifact rejection.

To test for systematic differences in articulatory movements between the *Imagery* and *Reading* conditions and, more critically, between the imagined syllables we conducted a decoding analysis. We used the same procedure described above (see *Temporal Generalization*). However, we only fitted each linear classifier model over the diagonal of the TG matrix (same training and testing time) since in this case we were not interested in the generalization across time. This decoding analysis was conducted at the single-subject level. Clusters of statistically significant decoding within subjects were determined using a cluster-based permutation test. For each timepoint (4ms) the classification procedure was repeated 1000 times after permuting the condition labels to build a null distribution. Scores falling beyond the 95^th^ percentile of the null distribution were then grouped into clusters (minimum cluster size = 10 samples). Next, the values inside the clusters were summed and used to compute a cluster statistic distribution. Clusters with *p* < 0.05 were considered statistically significant. Classification performance across subjects was evaluated using a statistical test for ROC AUC scores that accounts for the number of trials of each participant (Olivetti et al.; 2012). This was necessary to avoid statistical biases introduced by the uneven amounts of trials across participants resulting from the artefact rejection step. The statistical test was designed within a Bayesian hypothesis testing framework and, therefore, it returns a Bayes Factor as output.

## Notes

### Competing Interest Statement

The authors have declared no competing interest.

### Summary of Updates

The manuscript has been improved in terms of readability and clarity. To increase overall coherence, the WMC analysis (although potentially very interesting) has also been replaced by decoding analyses more in keeping with the rest of the paper.

