## Supplementary Information for "Decoding imagined speech reveals speech planning and production mechanisms"

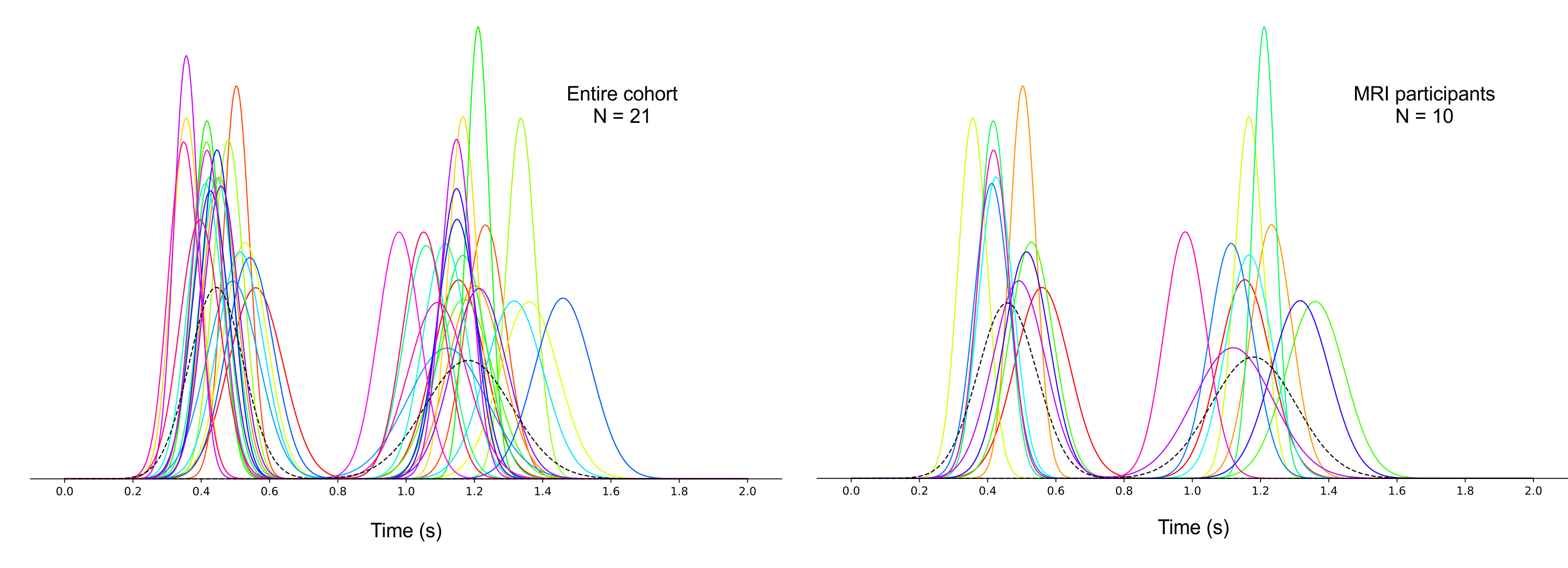

**Fig S1. Overt syllable onsets distributions.** Participants completed a minimum of 1 practice block on the day of the MEG acquisition with the exact same paradigm (Fig 1) but with *overt* productions of the syllables. Each production (2 syllables per trial) was annotated for syllable onset and coda. For each participant, we fitted a Gaussian curve on their distribution of syllable onsets for each production. The figure shows the fitted Gaussians of all participants (left) and of participants in the MRI sample (right). A Gaussian was also fitted to the overall distribution of participants’ onsets for each production in each group. These group-level Gaussians are plotted in dashed lines. Both groups exhibit a similar pattern with more precise productions for the first syllable and much more dispersion for the second, both intra- and inter-subject.

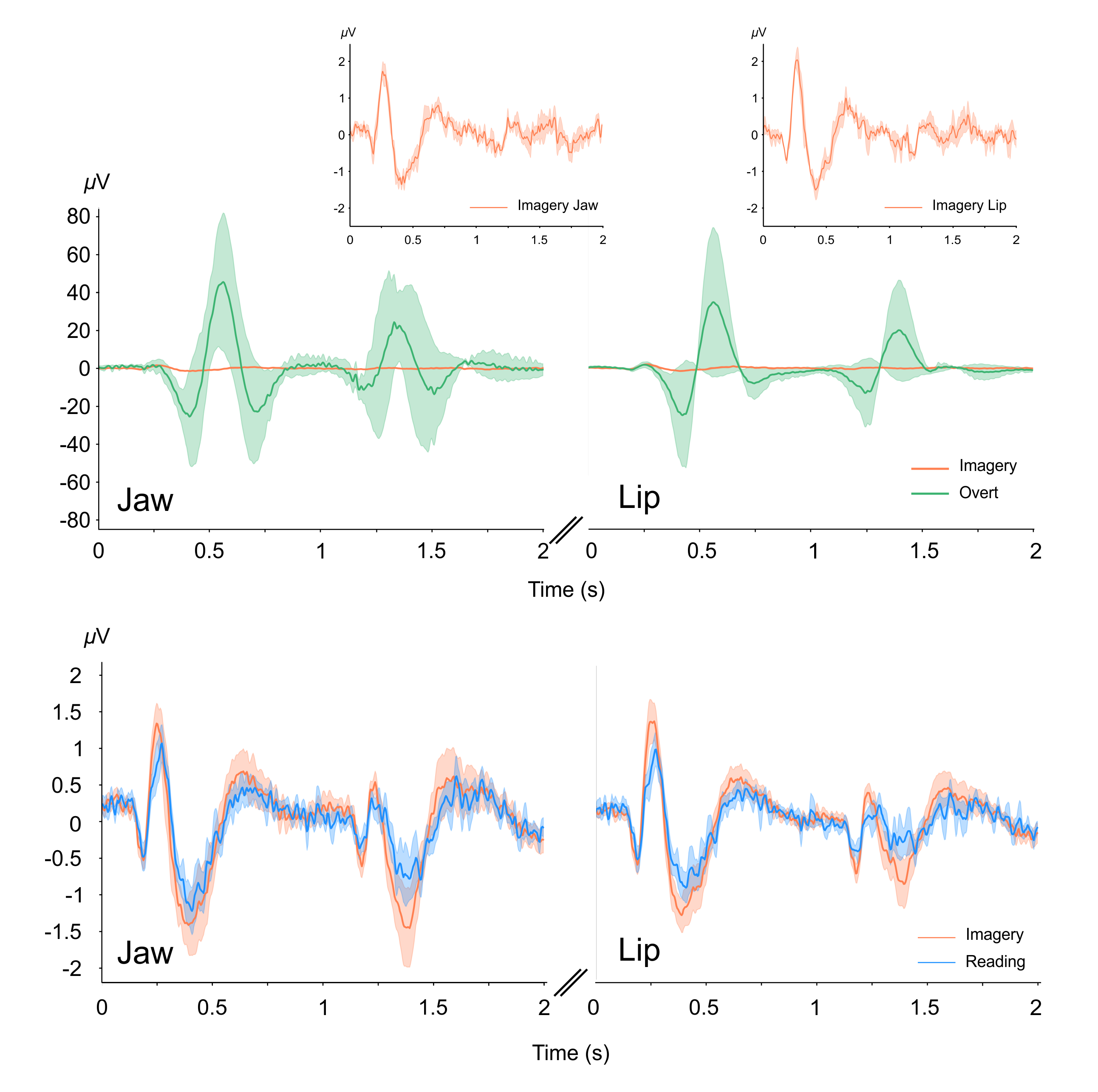

**Fig S2. Average EMG signal for all syllables (*pa*, *ta*, and *ka*). A. EMG signals during *Overt* and *Imagery* trials.** To put micromovements into context, we acquired EMG data from an additional 4 participants producing the three syllables (*pa*, *ta*, and *ka*). The plot shows their averaged activity for the jaw channel (left) and for the lip channel (right) (green lines with SEM) against the micromovement activity of the main cohort (N = 21; orange lines) during *Imagery*. The latter are plotted separately above. **B.** **EMG signal during *Reading* and *Imagery* trials (N = 21).** Micromovements were detected in both conditions, with subtle differences in amplitude between the two (Fig S5 below).

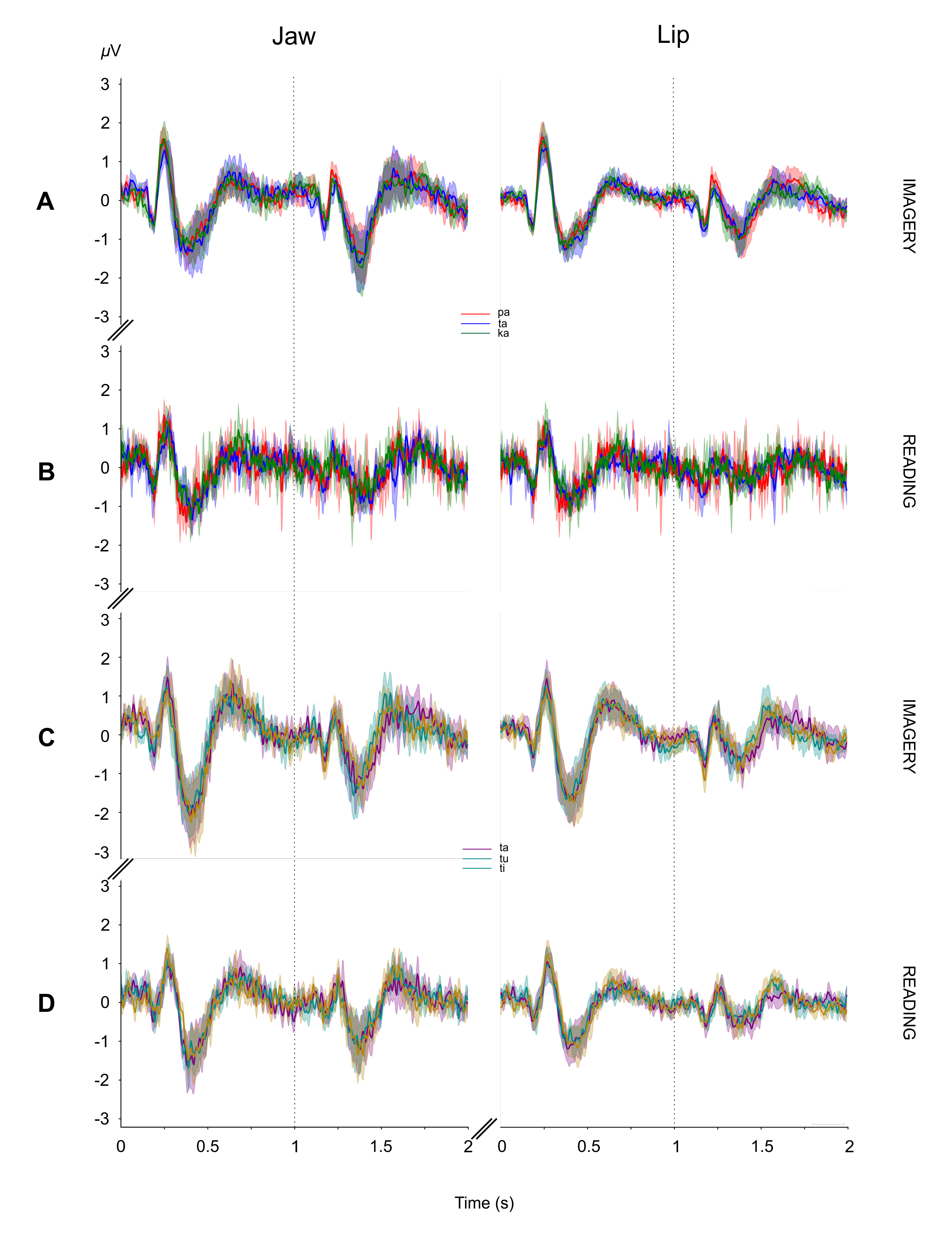

**Fig S3. Indistinguishable micromovements for syllables *pa*-*ta*-*ka* and *ta*-*tu*-*ti* both during *Imagery* and *Reading* trials.** Average EMG signals for the syllables *pa*-*ta*-*ka* during *Imagery* (**A**) and *Reading* (**B**) with SEM (N = 21)*.* Average EMG signals for the syllables *ta*-*tu*-*ti* during *Imagery* (**C**) and *Reading* (**D**) with SEM (N = 9)*.* Micromovements were almost identical for the 6 syllables employed.

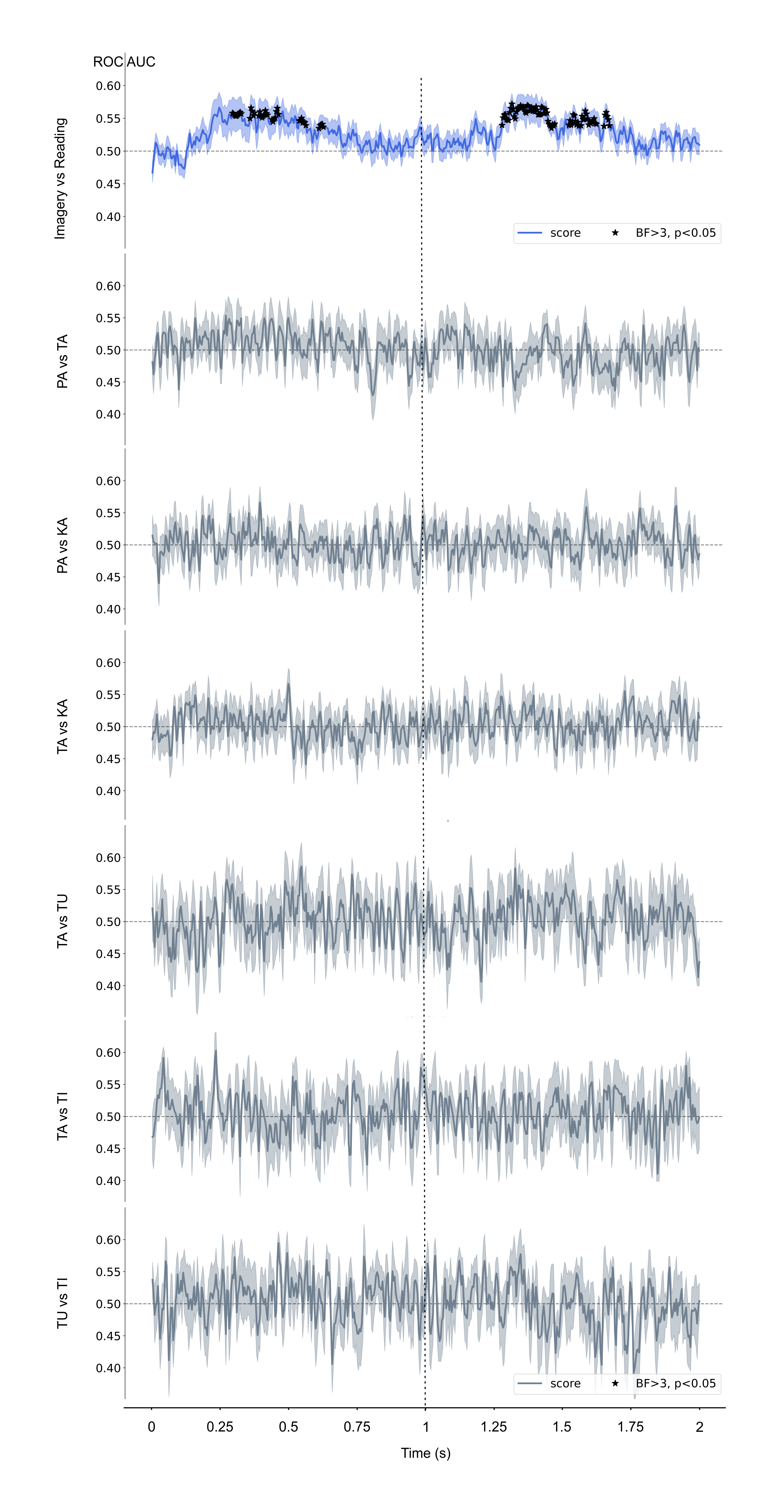

**Fig S4. Decoding using the EMG data.** The analyses of the EMG data show no significant decoding in any of the syllable contrasts despite the small differences between the main conditions (indicated by asterisks). Blue = ROC AUC scores and SEM for the *Imagery* vs *Reading* contrast. Grey lines = ROC AUC scores and SEM for the syllable contrasts (indicated on the left of each plot). Black asterisks indicate statistically significant decoding.

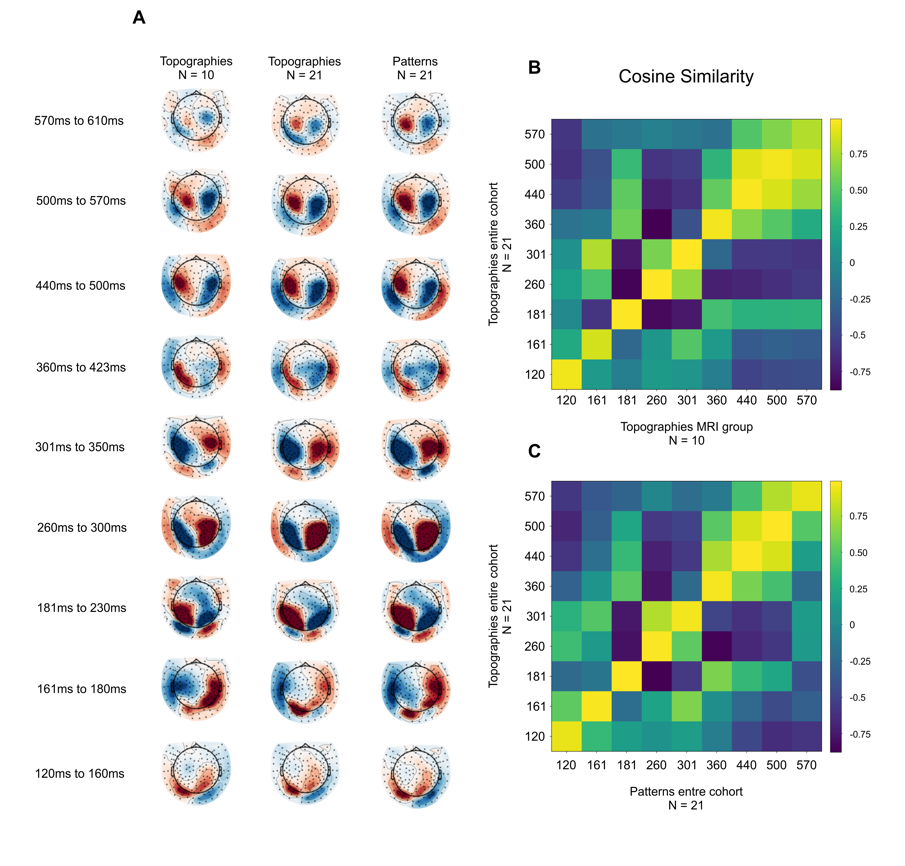

**Fig S5. Topographies for the MRI subgroup faithfully represent the neural sequence for imagery**. **A.** Average topographies across time for the *Imagery* condition for the MRI subgroup (left) and for the entire cohort (middle) alongside the syllable decoding classifier patterns (coefficients; right). **B.** Cosine similarity matrix comparing the topographies of the MRI subgroup (*y*-axis) with those of the entire cohort (*x*-axis). **C.** Cosine similarity matrix comparing the topographies of the entire cohort (*y*-axis) and syllable decoding patterns for the entire cohort (*x*-axis). The obvious similarities across time (high diagonal values) in all cases indicate that the MRI sample’s topographies for the *Imagery* condition faithfully represent the neural processes underlying speech imagery.

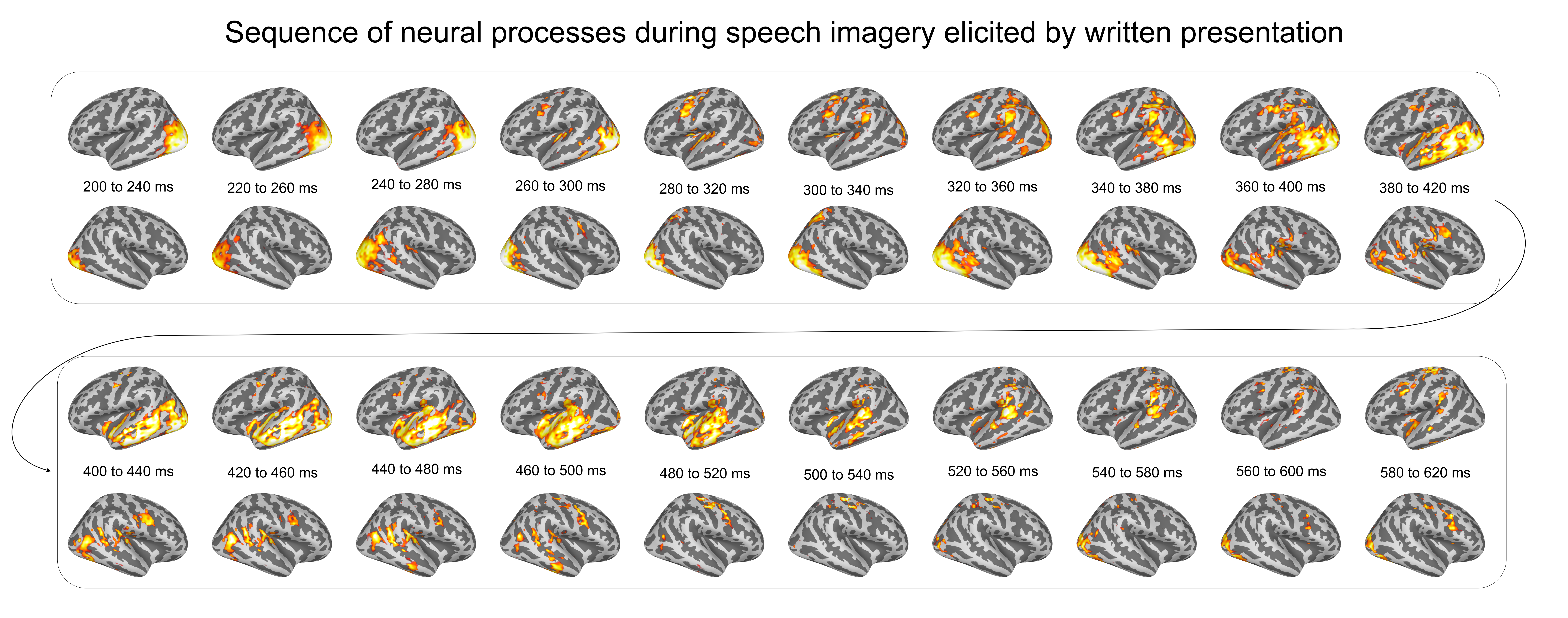

**Fig S6. Sequence of neural events during imagery event 1.** Average source space projections for the MRI group during imagery event 1 between 200 ms and 620 ms after syllable presentation. Each rendering shows the average of 20 ms of data. For display purposes, source space activity was thresholded at minima ranging between 1.94 and 6.24 and maxima between 2.42 and 8.49 units.

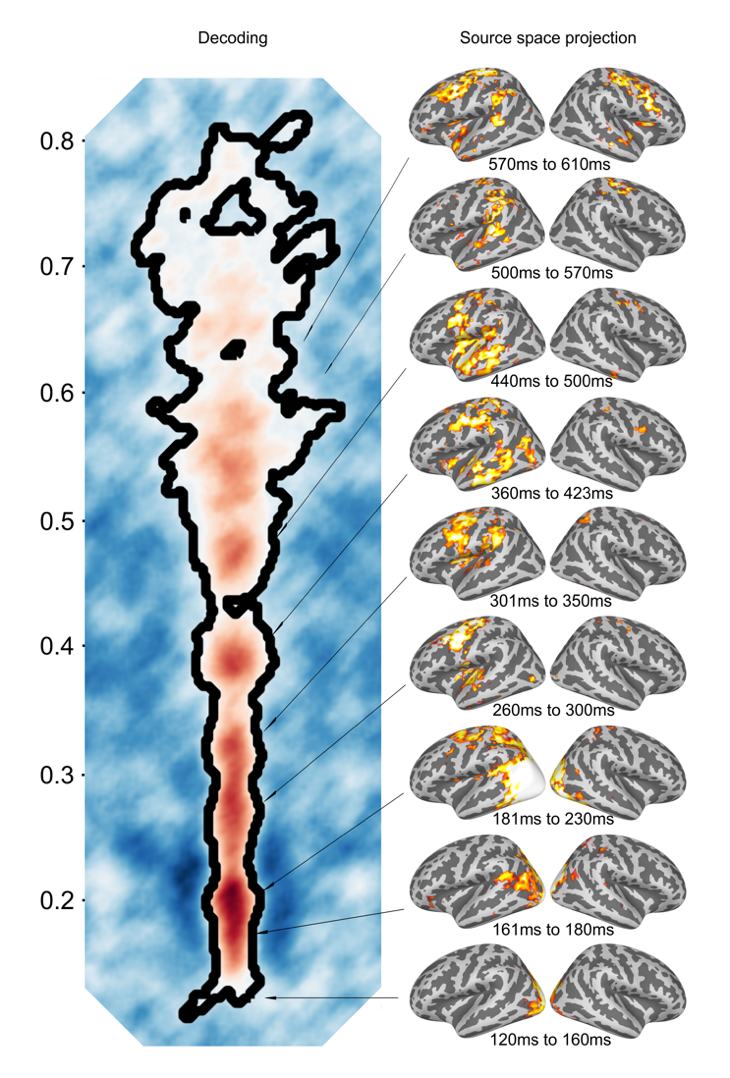

**Fig S7. Clusters of syllable decoding reveal a processing cascade during speech *Imagery (Imagery* minus *Reading*)*.*** Evoked activity estimated with sLORETA (ref; right panel; MRI group, N = 10) for *Imagery* minus *Reading* corresponding to the clusters of significant syllable decoding in the *Imagery* condition, event 1 (Fig. 3B). Spatial patterns were thresholded at +-40 fT. Evoked response topographies were thresholded at +-20 fT. Source space activity was thresholded at minima ranging between 0.6 and 1.61 and maxima between 1.14 and 2.59 units for display purposes.

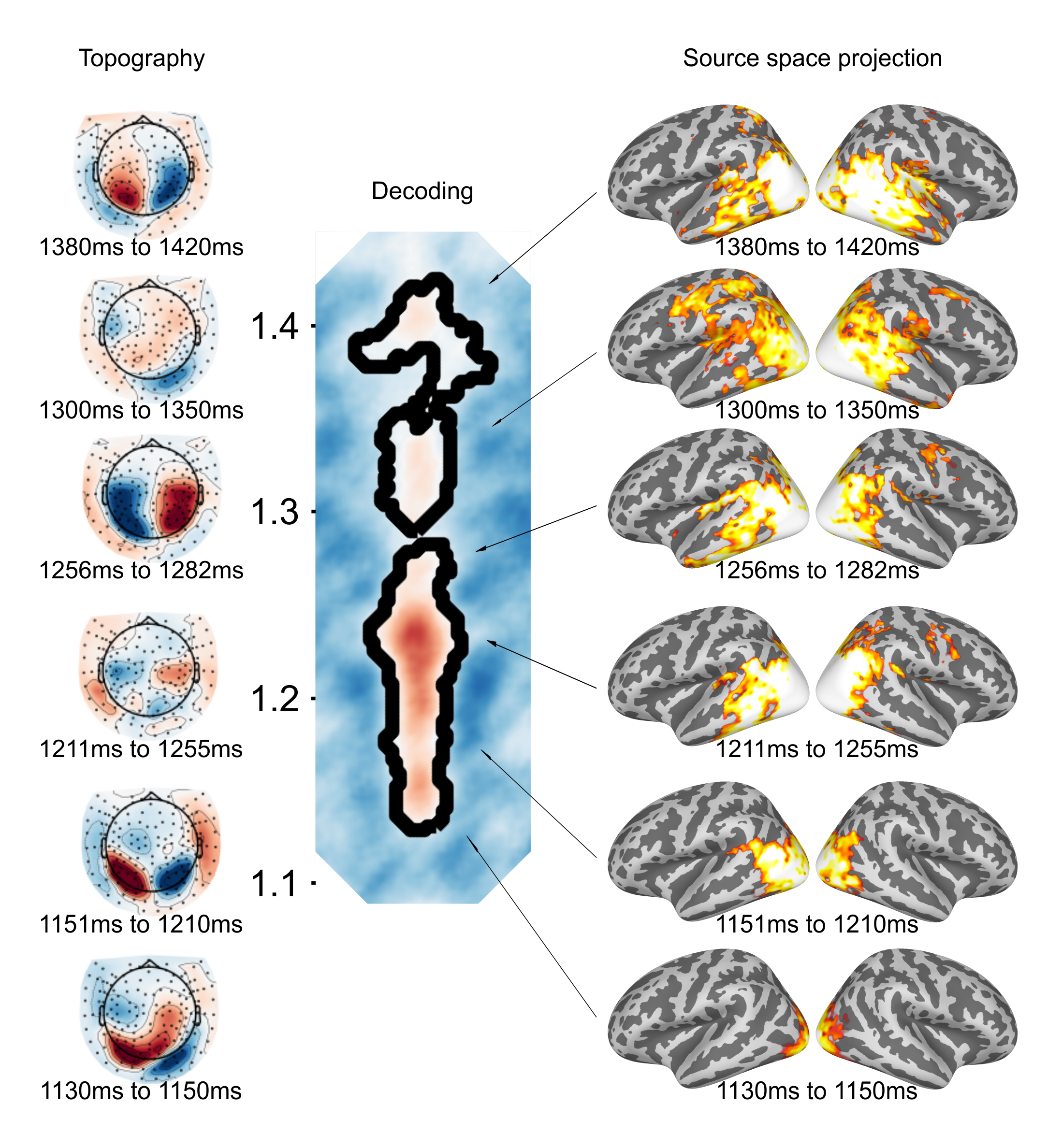

**Fig S8. Clusters of syllable decoding reveal a processing cascade during speech *Imagery (*event 2).** Evoked response topographies (*Imagery*) averaged over participants (left panel; N = 21) and evoked activity estimated with sLORETA (right panel; MRI group, N = 10) corresponding to the clusters of significant syllable decoding in the *Imagery* condition, event 2 (Fig. 2B). Evoked response topographies were thresholded at +-20 fT. Source space activity was thresholded at minima ranging between 1.53 and 3.42 and maxima between 1.92 and 4.58 units for display purposes.

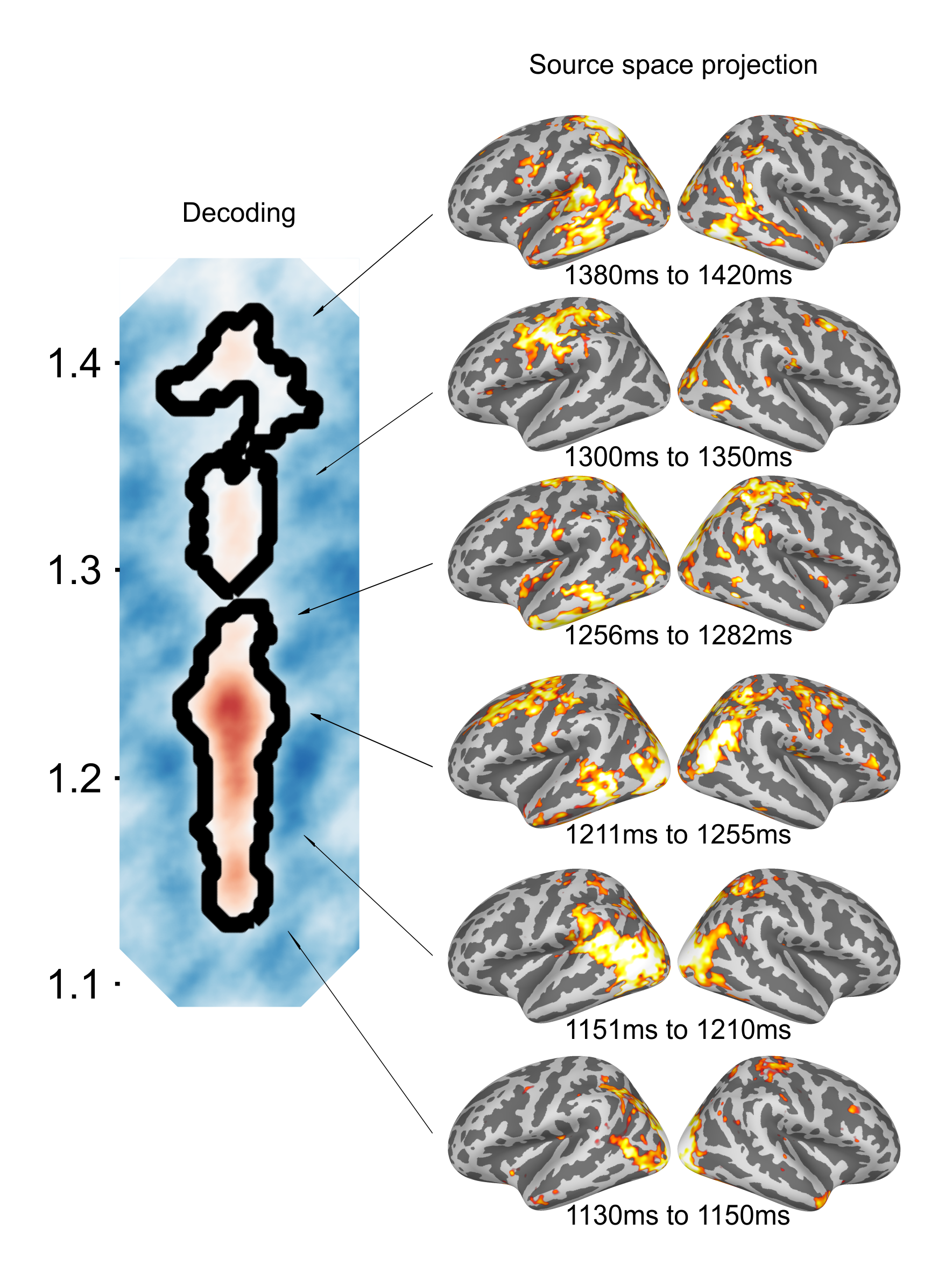

**Fig S9. Clusters of syllable decoding reveal a processing cascade during speech *Imagery,* event 2 *(Imagery* minus *Reading*).** Evoked activity estimated with sLORETA for *Imagery* minus *Reading* (right panel; MRI group, N = 10) corresponding to the clusters of significant syllable decoding in the *Imagery* condition, event 2 (Fig. 2B). Evoked response topographies were thresholded at +-20 fT. Source space activity was thresholded at minima ranging between 0.54 and 0.9 and maxima between 0.81 and 1.85 units for display purposes.

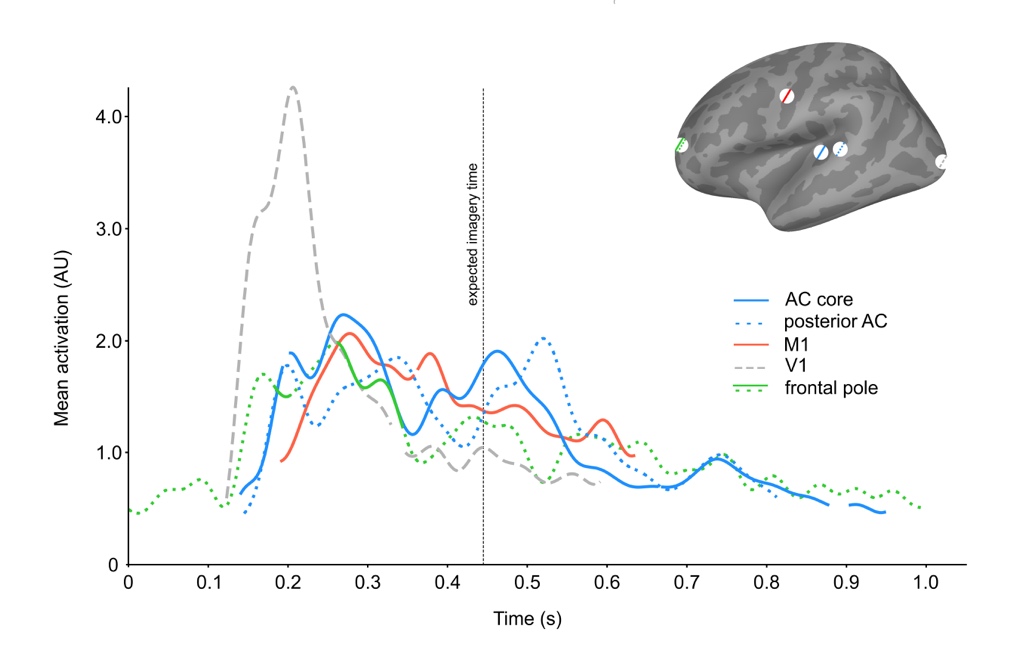

**Fig S10. Time courses of auditory and motor regions against two control regions during speech imagery.** The time courses of auditory (core and posterior) and motor regions are plotted along with two control regions (V1 and frontal pole). Although the time courses of motor and auditory regions interweave in line with SFC, the time courses of control regions show no apparent relationship. Time courses were z-scored and low-pass filtered at 20 Hz (double pass Butterworth of order 5) for display purposes only. Blank segments in the lines indicate non-significant times, except for the frontal pole, whose time course is shown in full with significant portions in solid green color.

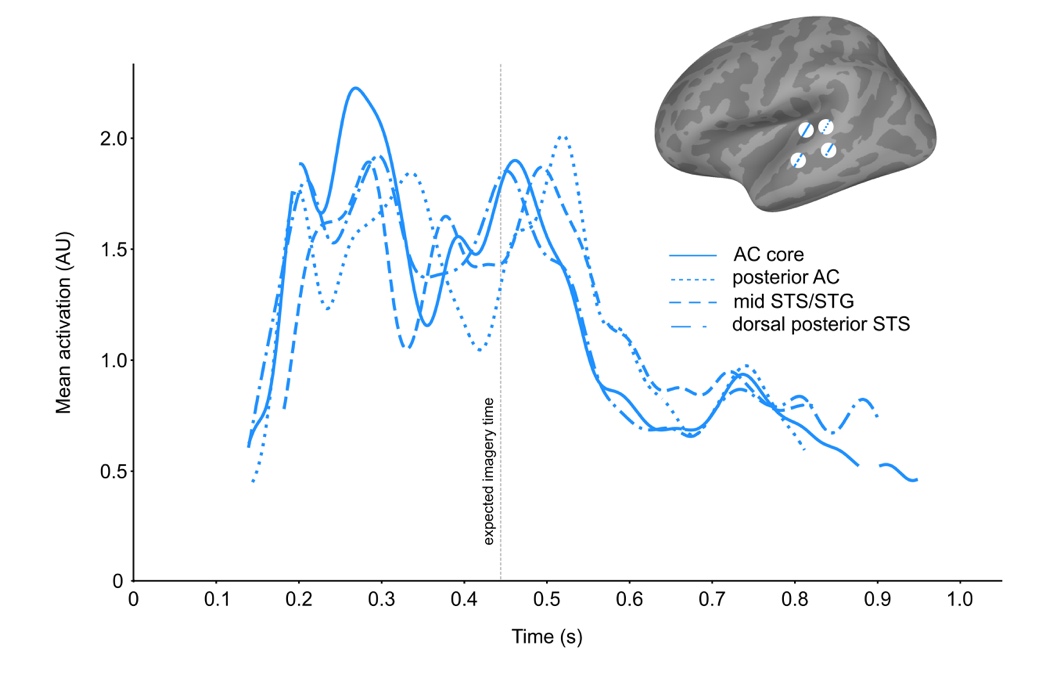

**Fig S11. Time course Auditory regions during speech imagery.** Widespread activity in auditory regions can be observed both before the expected time of imagery, reflecting motor preparation, and at the expected time of imagery, corresponding to the percept of the imagined syllable. Blank segments in the lines indicate non-significant times. Time courses were z-scored and low-pass filtered at 20 Hz (double pass Butterworth of order 5) for display purposes only.

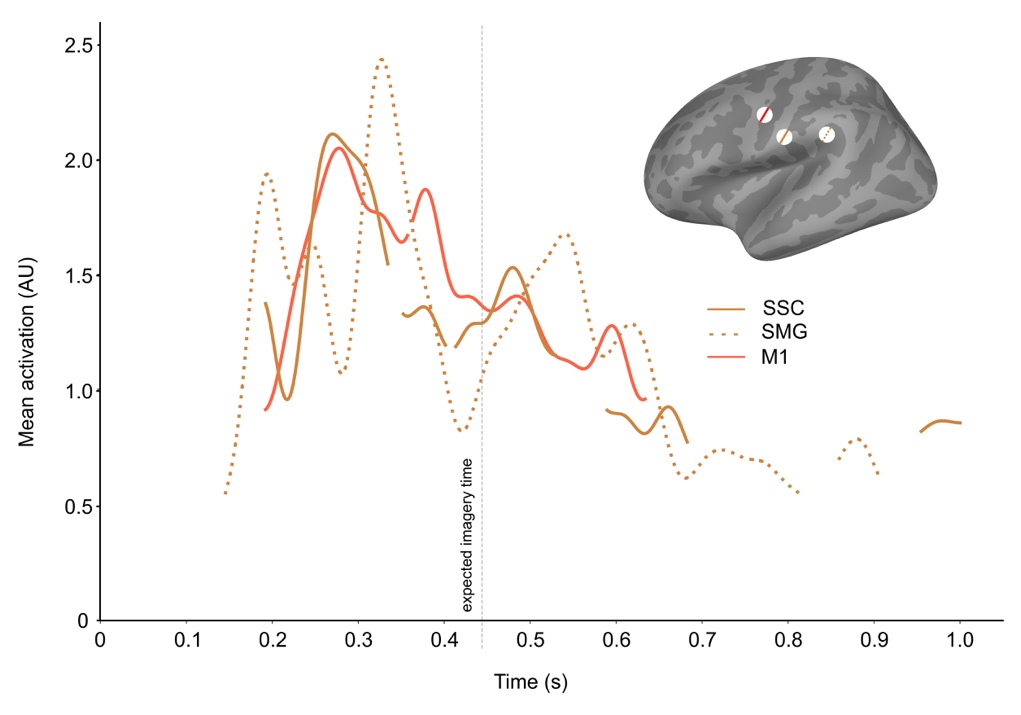

**Fig S12. Time course motor and somatosensory regions during speech imagery.** SFC theories hypothesize an internal feedback loop for motor planning and an external loop for post-production feedback, both characterized by feedforward and feedback processes between motor and sensory regions. The time courses of two ventral somatosensory cortex regions and motor regions show activity pre and post internal speech production in a sequence consistent with the hypothesized double feedback loop, paralleling the dynamics of auditory regions. Blank segments in the lines indicate non-significant times. Time courses were z-scored and low-pass filtered at 20 Hz (double pass Butterworth of order 5) for display purposes only. SSC = somatosensory cortex; SMG = supramarginal gyrus.

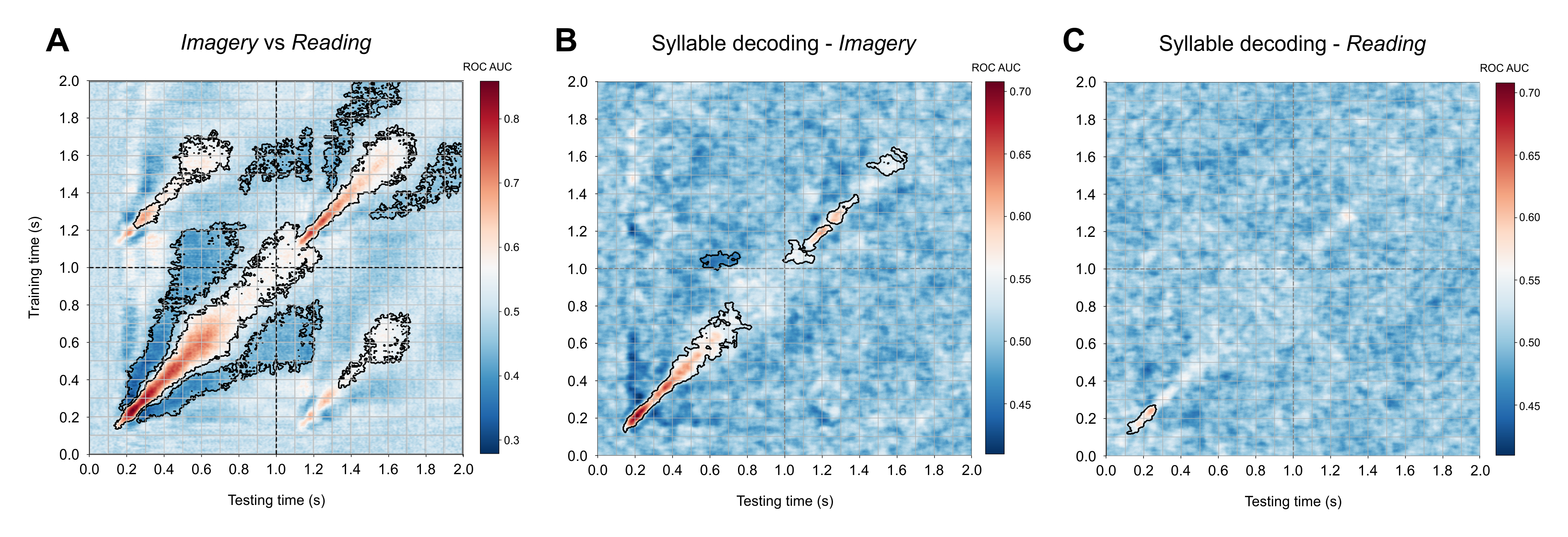

**Fig S13. Temporal generalization matrices replicating previous results (Fig 3) with different syllables (*ta*, *tu*, and *ti*). A.** Average TG matrix (N = 9) for the contrast *Imagery* vs *Reading*. ROC AUC = Receiver Operative Curve Area Under the Curve (chance = 0.5). **B.** and **C.** TG matrices for the pairwise contrasts between syllables (*ta* vs. *tu, ta* vs. *ti,* and *tu* vs. *ti)* for each condition (*Imagery* and *Reading*, respectively*)* first averaged within subject and then across subjects (N = 9). Clusters of statistically significant decoding (*p <* 0.05; black contour lines) were in all cases determined at the second level of analysis via a cluster-based permutation test across subjects (1000 permutations; two-tailed). Statistical significance indicates consistence across subjects, while high ROC AUC values reflect robust classifier performance on discriminating the contrasts. A great degree of similarity to previous results is observed, but for more robust and protracted decoding, particularly at the expected imagery time (event 1 median: ~349ms; event 2 median: ~1098ms).

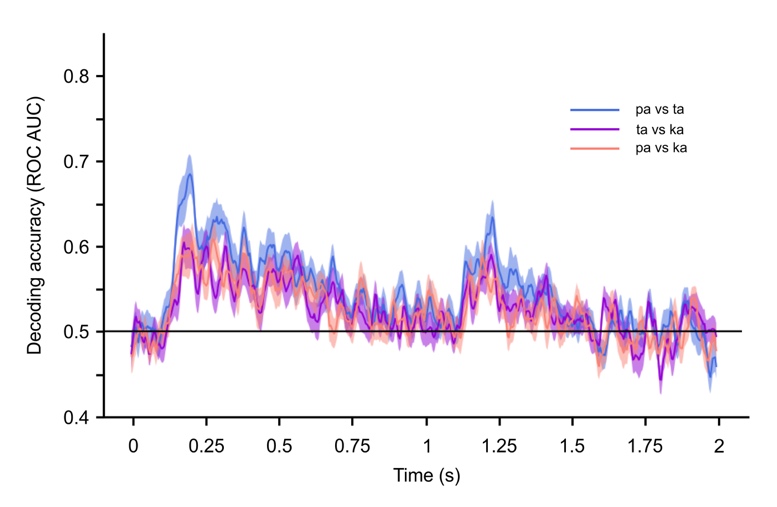

**Fig S14. Pairwise syllable decoding contrasts for the *pa*-*ta*-*ka* set** (/pa/ vs. /ta/, /pa/ vs. /ka/, and /ta/ vs. /ka/). Shaded regions show the standard error of the mean. Vertical dashed lines indicate the expected imagery onsets for each of the imagery events. Solid black line indicates chance level.

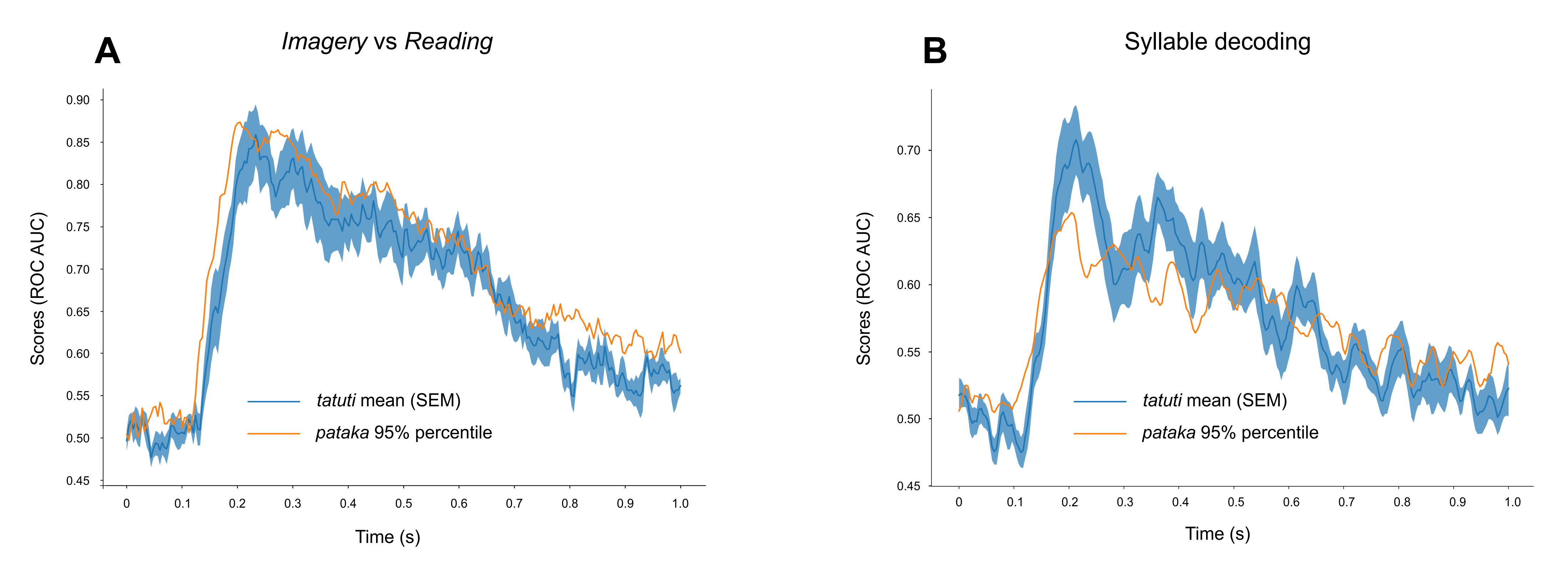

**Fig S15. Comparison of decoding scores for *pataka* vs *tatuti****.* **A.** Mean decoding scores across subjects (N = 9) and SEM for the *tatuti* set for the contrast between *Imagery* and *Reading* conditions against the 95^th^ percentile for the same contrast in the *pataka* set generated from ~300K permutations of 9 subjects from the larger cohort of 21 participants. **B.** Mean decoding scores across subjects (N = 9) and SEM for the *tatuti* set for the contrast between syllables (i.e., syllable decoding) against the 95^th^ percentile for the same contrasts in the *pataka* set generated from ~300K permutations of 9 subjects from the larger cohort of 21 participants. Significant differences in syllable decoding accuracy span large portions of event 1 between 185ms and 258ms and between 340ms and 451ms, suggesting that decoding is more robust on the vowel (acoustic) rather than the consonant (articulatory) representational space. ROC AUC = Receiver Operative Curve Area Under the Curve (chance = 0.5); SEM = Standard Error of the Mean.

**Table S1. Summary statistics for participants’ overt syllable productions.** Median onsets represent the expected time of imagery for the different cohorts.

|  | Syllable 1 | | Syllable 2 | | Syllable 1 | | Syllable 2 | | Transition 1 | | Transition 2 | |
| --- | --- | --- | --- | --- | --- | --- | --- | --- | --- | --- | --- | --- |
|  | Median onset (ms) | IQR (ms) | Median onset (ms) | IQR (ms) | Median duration (ms) | IQR (ms) | Median duration (ms) | IQR (ms) | Median duration (ms) | IQR (ms) | Median duration (ms) | IQR (ms) |
| *pa-ta-ka* (N = 21) | 436 | 99 | 1175 | 146 | 194 | 59 | 192 | 51 | 114 | 24 | 116 | 24 |
| *pa-ta-ka* (N = 10) | 444 | 116 | 1185 | 140 | 205 | 55 | 199 | 50 | 111 | 23 | 111 | 24 |
| *ta-tu-ti* (N = 9) | 349 | 90 | 1098 | 127 | 195 | 43 | 199 | 42 |  |  |  |  |
